# Unraveling synergistic and antagonistic effects of simultaneous versus single hypoxia-salt conditions in an evolutionary adapted plant species

**DOI:** 10.1101/2025.10.09.681434

**Authors:** Angelina Jordine, Julia Alt, Joost T. van Dongen, Lisa Fürtauer

## Abstract

Plants frequently encounter simultaneous stressors, requiring complex adaptive responses across regulatory levels. Although morphological and metabolic effects of combined hypoxia-salt stress are known in halophytes, transcriptional impacts remain largely unexplored. This study uses RNA sequencing to analyze *Salicornia europaea* which is an intertidal salt-marsh plant naturally adapted to these conditions. Combined hypoxia-salt stress induced a distinct gene expression profile in which 16% of genes were exclusively changed. Synergistic, antagonistic, and additive effects were observed across all evaluated functional pathway categories. A data-driven analysis of carbohydrate metabolism, cellular respiration/fermentation, and amino acid pathways revealed that antagonistic effects were more prevalent than synergistic ones in both roots and shoots. Notably higher gene expression levels during hypoxia-salt of sucrose biosynthesis (consistent with salt), *sucrose synthase* (*SUS*) and *trehalose-6-phosphate phosphatase* (*TPP*, consistent with hypoxia) indicate enhanced sucrose and trehalose metabolism. The parallel down-regulation of invertase genes (consistent with hypoxia) suggests strategic carbon flux redistribution for optimized energy supply. Under hypoxic conditions, *lactate dehydrogenase* (*LDH*) expression was up-regulated, indicating active lactate fermentation rather than ethanol production via *alcohol dehydrogenase* (*ADH*). Enhanced proline synthesis under combined stress suggests improved osmoprotection. These insights into transcriptional reprogramming under single and combined hypoxia-salt conditions emphasize the intricate regulatory strategies plants utilize to manage concurrent stressors, showcasing their ability to adapt and develop stress resilience.

## Introduction

Plants established themselves in all terrestrial ecosystems by adapting to particular environmental conditions and developing tolerance mechanisms (Waters, 2003). These mechanisms allow plants to survive an thrive in extreme habitats, like hot and cold deserts, alpine regions and coastal areas (Billings and Mooney, 1968; Bliss, 1971; Gutterman, 2012). Consequently, in natural habitats, plants are often exposed to multiple abiotic stress factors simultaneously rather than being exposed to a single one. Survival in such conditions necessitates complex adaptation and tolerance mechanisms that include genome adaptations and integrate signaling transduction networks, transcriptional as well as metabolic adaptations upon abiotic stress (Behr et al., 2017; Bulut et al., 2025; Garg et al., 2013; Zandalinas et al., 2021). Consequently, research on simultaneous stress factors in various combinations has increased rapidly in recent years (Balfagón et al., 2019; Bulut et al., 2025; Fürtauer et al., 2018; Jordine et al., 2024; Lu et al., 2019; Mittler, 2006; Rizhsky et al., 2004; Sinha et al., 2024; Zandalinas and Mittler, 2022; Zandalinas et al., 2021).

Examinations of combinations of drought, heat, high light and salinity stress simultaneously suggested that the adaptive responses to these combined factors cannot be directly predicted from how plants respond to each individual condition (Mahalingam, 2015; Mahalingam et al., 2021; Rizhsky et al., 2004). A “dominant stressor” may dictate the combined response based on severity (Pandey et al., 2015). Simultaneous stresses lead to complex interactions within their metabolic responses and reprogramming processes, involving intricate regulatory strategies such as signaling events that lead to transcriptional regulation and modified enzyme activities. Being exposed to multiple stressors, can lead to an overall additive, synergistic or antagonistic effect for the plant. Synergy occurs when combined stressors have a greater effect than the sum of their individual impacts. In contrast, antagonistic interactions reduce the overall effect compared to the sum of individual stressors, while additive effects result in an impact equal to their sum (Alptekin and Kunkowska, 2024; Zandalinas and Mittler, 2022).

Synergistic, antagonistic, and additive stress response concepts have mainly been used to describe the overall effect of simultaneous stress factors, but are seldom applied to metabolic or gene expression changes (Alptekin and Kunkowska, 2024; Zandalinas and Mittler, 2022). Hypoxia and salinity stress often co-occur in coastal regions, where plants are naturally exposed to fluctuating environmental conditions. In these dynamic habitats, especially in salt marshes, tides and freshwater input *via* rainfall create frequent changes in oxygen availability and salinity levels (Glup, 1985). While the combination of hypoxia and salt stress has been studied at the morphological and metabolite levels in tolerant plant species such as *Salicornia europaea* (Jordine et al., 2024), *Suaeda maritima* (Behr et al., 2017), *Phragmatis australis* (Gorai et al., 2010) and *Beta vulgaris* (Behr et al., 2021), this study focuses explicitly on transcriptional reprogramming under single and combined hypoxia-salt conditions in *Salicornia europaea*.

Hypoxia and salinity independently trigger distinct metabolic responses in plants, affecting fundamental processes such as i) carbohydrate metabolism, ii) amino acid (AA) metabolism and iii) respiration and energy production. For example, cellular respiration is generally reduced under low oxygen conditions (Zabalza et al., 2009). During hypoxia, transcriptional reprogramming occurs in parts of the carbohydrate metabolism (trehalose metabolism) that controls the sugar influx into glycolysis (Barding et al., 2013; Liu et al., 2005). Glycolysis itself and fermentation are also reprogrammed resulting in anaerobic metabolism (Licausi, 2012; Liu et al., 2005). Additionally also amino acid metabolism, ethylene signaling and nitrogen usage are significantly altered under hypoxia (Barding et al., 2013; Liu et al., 2005). All of these adaptive mechanisms are focused on energy conservation to prevent the plant from energy shortage (Cho et al., 2021; Sasidharan and Mustroph, 2011).

Salt stress on the other hand, leads to ionic imbalance and osmotic stress, disturbances in cellular homeostasis, variations in respiration rates and energy production, generation of reactive oxygen species (ROS) and reduced water availability (O’Leary et al., 2019; Shahid et al., 2020; Yang and Guo, 2017). For example, some respiratory pathways like the tricarboxylic acid (TCA) cycle or mitochondrial electron transport chain (mETC) use amino acid degradation products as alternative substrates among others. Amino acid degradation products serve as alternative substrates that feed into carbohydrate metabolism, supporting energy homeostasis and recovery under salt stress (Heinemann and Hildebrandt, 2021; Zhang et al., 2017).

Under hypoxic conditions, energy production *via* aerobic respiration is reduced, causing a metabolic shift toward anaerobic fermentation. This change leads to enhanced sugar consumption, mobilization, and transport to maintain energy supply. However, sucrose synthesis is down-regulated, resulting in no net accumulation of sugars (Cho et al., 2021; van Veen et al., 2024). In contrast, under salt stress, adenosine triphosphate (ATP) production remains largely unaffected, but energy demand increases due to the activation of stress-related processes. Despite a general inhibition of sugar consumption, mobilization, and transport, sucrose synthesis itself is enhanced. This leads to the accumulation of soluble sugars, acting mainly as osmoprotectants against osmotic imbalance from salinity (Sellami et al., 2019).

Moreover, contrasting responses for individual hypoxia and salt stress are reported for amino acid metabolism. Glutamate (Glu) is degraded *via* GLUTAMATE DEHYDROGENASE (GDH) to recycle ammonium and conserve energy during hypoxia. Additionally, the conversion of pyruvate to alanine (Ala) is a key mechanism for energy-efficient nitrogen recycling (Diab and Limami, 2016). In contrast, salt stress suppresses alanine metabolism, as indicated by reduced levels of Ala, asparagine (Asn), and glutamine (Gln) (Sellami et al., 2019). Furthermore, GLUTAMINE SYNTHETASE activity is up-regulated in roots under salt stress to support ammonium assimilation (Teixeira and Fidalgo, 2009). Proline strongly accumulates as an osmoprotectant, and this response is supported by the salt-induced activation of amino acid transporters (e.g. AMINO ACID PERMEASE 1, AAP1), which facilitate proline uptake from external sources (Misra and Gupta, 2005; T. Wang et al., 2017). In contrast, under hypoxic conditions, proline does not accumulate, and no specific transporters for its uptake are induced. Under oxygen deficiency, respiration in the root tissue of *Pisum sativum* (pea) decreases, while salt stress generally either increases respiration or has no effect, depending on the species (Jacoby et al., 2011; Zabalza et al., 2009).

Combined hypoxia-salt stress lead to increased proline accumulation, alterations in carbohydrate levels, elevated osmotically relevant metabolites, and modifications in the tricarboxylic acid (TCA) cycle, indicating significant metabolic adaptation (Behr et al., 2017; Behr et al., 2021; Gorai et al., 2010). Although these studies provide important insights into the physiological response to simultaneous hypoxia and salt stress, they have only been a limited number of studies on the transcriptional level (Garg et al., 2013). The investigation of combined salt-hypoxia conditions using a naturally tolerant species like *Salicornia europaea*, which occurs in salt coastal marshes, can reveal insights and enhance our understanding of these conditions. This species serves as an ideal model plant for studying adaptive mechanisms to combined hypoxia and salt stress, as demonstrated by our previous research on selected transcriptional responses under simultaneous salinity and flooding, as well as sequential salinity and hypoxia (Jordine et al., 2024). Notably, the expression patterns of hypoxia-responsive genes were altered under sequential salinity and hypoxia compared to individual stress treatments. These findings underscore the potential of *Salicornia europaea* as a valuable model to dissect gene regulatory mechanisms under combined hypoxia and salinity stress (Jordine et al., 2024).

In the present study, we aimed to identify hypoxia-salt specific gene expression changes in hypoxia or salt related pathways in the adapted plant *Salicornia europaea* to draw physiological conclusions regarding simultaneous stress interactions. We elucidated the systemic hypoxia-salt specific gene expression changes on a larger scale *via* RNA sequencing during single and simultaneous stress in both shoot and root. Utilizing a comprehensive data analysis approach, we demonstrate that simultaneous hypoxia-salt stress induced an enhanced unique stress response. This unique reaction mainly involved altered expression patterns in already known hypoxia or salt responsive pathways. Thereby, novel additive, synergistic and antagonistic regulations were found. Driven by data, we focused on i) carbohydrate metabolism, ii) cellular respiration/fermentation, and iii) amino acid pathways to compare gene expression level changes and determined individual antagonistic, synergistic and additive effects.

## Material and Methods

### Plant Material and Cultivation

*Salicornia europaea* seeds were obtained from a seed distributer (Rühlemanńs Kräuter und Duftpflanzen, Horstedt, Germany). The cultivation was conducted in a growth chamber under controlled conditions with 16 h light (290 µmol photons m^-2^ s^-1^) at 22°C. Seeds were sown on square plates with wet filter paper (Suppl. Fig. S1A) for germination. After two weeks, the seedlings were transferred into 1.5 ml reaction tubes, half filled with solid ½ Hoagland medium (2.5 mM KNO_3_, 2.5 mM Ca(NO_3_)_4_, 0.5 mM KH_2_PO_4_, 0.5 mM MgSO_4_, 50 µM KCl, 25µM H_3_BO_3_, 2.25 µM MnCl_2_, 1.9 µM ZnSO_4_, 1.5 µM CuSO_4_, 0.05 µM (NH_4_)_6_Mo_7_O_24_, 40 µM Fe-EDTA and 0.5% (w/v) Agarose) and pre-cultivated floating in liquid, aerated 1/2 Hoagland medium (Fig. S1B). After three weeks, plants were transferred to 50 ml reaction tubes and acclimated for 1 week in hydroponic culture containing liquid 1/2 Hoagland medium (main culture, Fig. S1C). Salinity treatment started with 6 week old plants.

### Salinity and Hypoxia Treatment

First, six-week-old *S. europaea* plants were acclimated to salt. Therefore, continuously 100 ml NaCl (3.5 M) was added to the hydroponic culture (culture volume: 7 L) per day over a total time of 12 days to a final concentration of 500 mM. Added NaCl-solution corresponded to the evaporation of Hoagland medium in the hydroponic culture. The continuous addition was chosen to avoid a salt shock reaction (Shavrukov, 2012). Two days after reaching the final NaCl concentration (8 week old plants), the hypoxia treatment (1% O_2_ (v/v)) was performed where plants were kept dark to avoid photosynthetic O_2_ generation during hypoxic conditions. The starting time point was the middle of the day (8 hours of light). After 2 h of hypoxia treatment the shoots and roots were immediately frozen into liquid nitrogen, and stored at -80°C. Further, plants only exposed to salt or hypoxia were harvested as well as control plants not subjected to salt or hypoxia at the same time point. For every condition (control (**C**), salt (**S**), hypoxia (**H**) and hypoxia-salt (**HS**)) four samples of shoots and roots, consisting of each three plants were harvested.

### RNA Sequencing

The frozen plant material was ground in liquid nitrogen and RNA was isolated as described previously (Jordine et al., 2024). Total RNA was quantified with the RNA Broad Range Assay Kits on an Invitrogen Qubit 4 Fluorometer (Thermo Fisher scientific Inc, Germany), according to manufacturer’s instructions. Additionally, the integrity of the RNA (RIN value *>* 8) was certified in an Agilent 2100 Bioanalyzer (Agilent Technologies Inc., Germany) using the RNA 6000 Nano Kit. For sequencing the RNA concentration was set to 40 ng/µl. Paired end sequencing was performed by Eurofins Genomics (Eurofins Genomics GmbH, Germany) including strand-specific cDNA library preparation and Illumina sequencing on the NovaSeq platform (2 x 150 sequence mode). The raw RNA-Seq data sets from this study can be accessed through NCBI under Bioproject ID PRJNA1256208 for shoot data and PRJNA1256210 for root data.

### Raw RNA Sequence Processing

Raw sequencing data was processed *via* the Anaconda v23.11.0 environment (Supplemental S2, Anaconda, 2023). Quality control of the raw reads was performed with FastQC v0.12.1 (Andrews, 2010), then each sample was cleaned from low-quality reads and adapter fragments (Trimmomatic v0.39; Bolger et al., 2014). The alignment to reference transcripts (Jordine et al., 2024) was performed with HISAT2 v2.2.1 (Kim et al., 2019). SAM files were converted into BAM files (SAMtools v1.13; Danecek et al., 2021) and featureCounts v2.0.1 was used to summarize the counts of all samples into one read count table (Liao et al., 2014). All downstream analysis were performed in RStudio v4.3.3 (R Core Team, 2017). A schematic depiction of further analysis steps can be found in supplemental material S3, with detailed session information in the supplemental S4. For further analysis the read count table was filtered for genes with reads in at least 3 out of the 4 replicates.

### Sample Similarity and Differential Analysis

The differences between RNA sequencing samples are primary driven by the genes with the highest counts, as they exhibit the most significant absolute differences between samples. Therefore regularized-logarithm transformation (rlog transformation, DESeq2 v1.42.1 Love et al., 2014) was applied on the reads to analyze the sample similarity. A principal component analysis from the individual shoot and root data set and the combined data set was performed. For the differential expression between the conditions, counts were normalized using the median of ratios normalization (DESeq2 v1.42.1). Pairwise-comparison was then conducted with the control samples (normoxia without salt) as reference. The p-values were Benjamini-Hochberg corrected Benjamini and Hochberg, 2000 with a statistical cutoff for false discovery rate of*α* = 0.05. Subsequently the log2FoldChange (log2FC) of genes with high dispersion was corrected using apeglm (Zhu et al., 2018). Finally, a gene was classified as differentially expressed gene (DEG) if the adjusted p-value was *<* 0.01 (significant DEG, sDEG). The Venn analysis (VennDiagramm v1.7.3) was conducted based on sDEG sets from salt, hypoxia and hypoxia-salt with additionally abs(log2FoldChange)*>*2. All sDEGs identified through variance and Venn analyses were merged into a gene list (Suppl. Table 1). This combined list served as the basis for a non-targeted analysis aimed at uncovering unknown patterns and pathways.

### Gene Categorization and Pathway Analysis

Functional categorization of the genes was conducted with Mercator4 v2.0 (Lohse et al., 2013). Afterwards, MapMan was used for pathway analysis and visualization of DEGs (Schwacke et al., 2019). A Wilcoxon rank-sum test, followed by Benjamini–Hochberg correction (Benjamini and Hochberg, 2000) for p-values, was applied to determine whether the combined expression values in a specific functional BIN, which describe biological contexts, differ significantly from the expression changes observed in the collection of genes from all other BINs.

### Analysis of Additive, Synergistic and Antagonistic Effects

To determine the additive effect for each gene, we analyzed the relationship between the observed log2FoldChanges under simultaneous hypoxia-salt (**HS**) and the combined effect of salt stress (**S**) and hypoxia (**H**). First, log2FoldChange values were converted into linear FoldChanges (*FC* = 2^log2FC^ and *FC* = (−1)·2^abs(-log2FC)^) and the expression changes in percent (*pEX*) were calculated, by adding one to negative FoldChanges or subtracting one from positive FoldChanges (*pEX* := −*FC* + 1 and *pEX* := +*FC* − 1). To quantify the additive effect of **S** and **H**, the percentages of these conditions were summed (*pEX***_S+H_** = *pEX***_S_** + *pEX***_H_**). Genes were classified based on their percentage **HS** change relative to the sum of **S** and **H** percentages as i) additive if within ±0.5 of this sum: *pEX***_HS_** ∈ [*pEX***_H+S_** − 0.5, *pEX***_H+S_** + 0.5] ii) synergistic if greater than this range: *pEX***_HS_** *> pEX***_H+S_** + 0.5 or iii) antagonistic if smaller than this range: *pEX***_HS_** *< pEX***_H+S_** − 0.5. For visualization the percentage changes were transformed back to linear FoldChange (*FC* = *pEX***_S+H_** ±1) and log2FoldChange (*log*2*FoldChange* = *log*2(*FC*)). Log2FoldChange of **HS** was plotted against the log2FoldChange of the sum of **S** and **H**, synergistic and antagonistic genes were highlighted. We selected for initial analysis known hypoxia or salt-responsive pathways (amino acid metabolism, carbohydrate metabolism and cellular respiration). In addition to the pathways known to be involved, all pathways with a high probability of involvement under at least one of the conditions were also examined. Subsequently, a list of all synergistic and antagonistic genes was created from this analysis.

## Results

### Gene Expression Patterns from Single and Simultaneous Hypoxia-Salt Stress Cluster Distinctly

For investigation of the fast transcriptional effects of *Salicornia europaea* during simultaneous 2-hour hypoxia combined with/without salt-adaptation, we conducted an RNA sequencing analysis on shoot and root material. RNA sequencing yielded 28-34 million cleaned reads per shoot sample, which were aligned to the reference transcriptome (Jordine et al., 2024) with an alignment rate of 63-69%. For root samples, RNA sequencing resulted in 17-26 million cleaned reads and a slightly reduced alignment rate of about 54-63% (Suppl. Fig. S5). The data set resulted in a total of 13086 shoot and 13231 root genes with open reading frames, excluding isoforms (Suppl. S6A). Out of these, we successfully annotated 7969 shoot and 8062 root genes (Suppl. Fig. S6B).

Principal Component Analysis (PCA) of annotated genes revealed four distinct clusters based on conditions for both shoots (Fig. 1A) and roots (Fig. 1B). In both shoot and root, a clear separation of the stress conditions was determined, with a variance explained for shoot of ∼58% and root ∼61% in principal components 1 (PC1) and 2 (PC2). In both PC1 (*>*37%), plants treated with salt (red-triangles and blue-squares) were clearly separated from non-salt-treated ones (green-diamonds and yellow-cycles). In both PC2 (*>* 20%) normoxic and hypoxic conditions were separated. This separation persisted when non-annotated genes were included in the analysis (Suppl. Fig. S7). For all four conditions, only a small variation was detected between the biological replicates. Combined analysis of shoot and root data showed clear separation by PC1 (*>* 54%), distinguishing root data (empty shapes) from shoot data (filled shapes) (Fig. 1C left, PC1-PC2) and PC2 separated again salt treatment. Additionally, analysis of PC2 and PC3 (24%, Fig. 1C right) clustered the conditions similar to single data sets. Overall, ∼78% of the variance could be explained by PC1, PC2 and PC3 in this combined analysis. These findings demonstrate that the combination of two stress factors lead to a distinct gene expression pattern compared to single stress treatments, with clear separation observed not only between stress conditions but also between plant organs.

**Figure 1:**
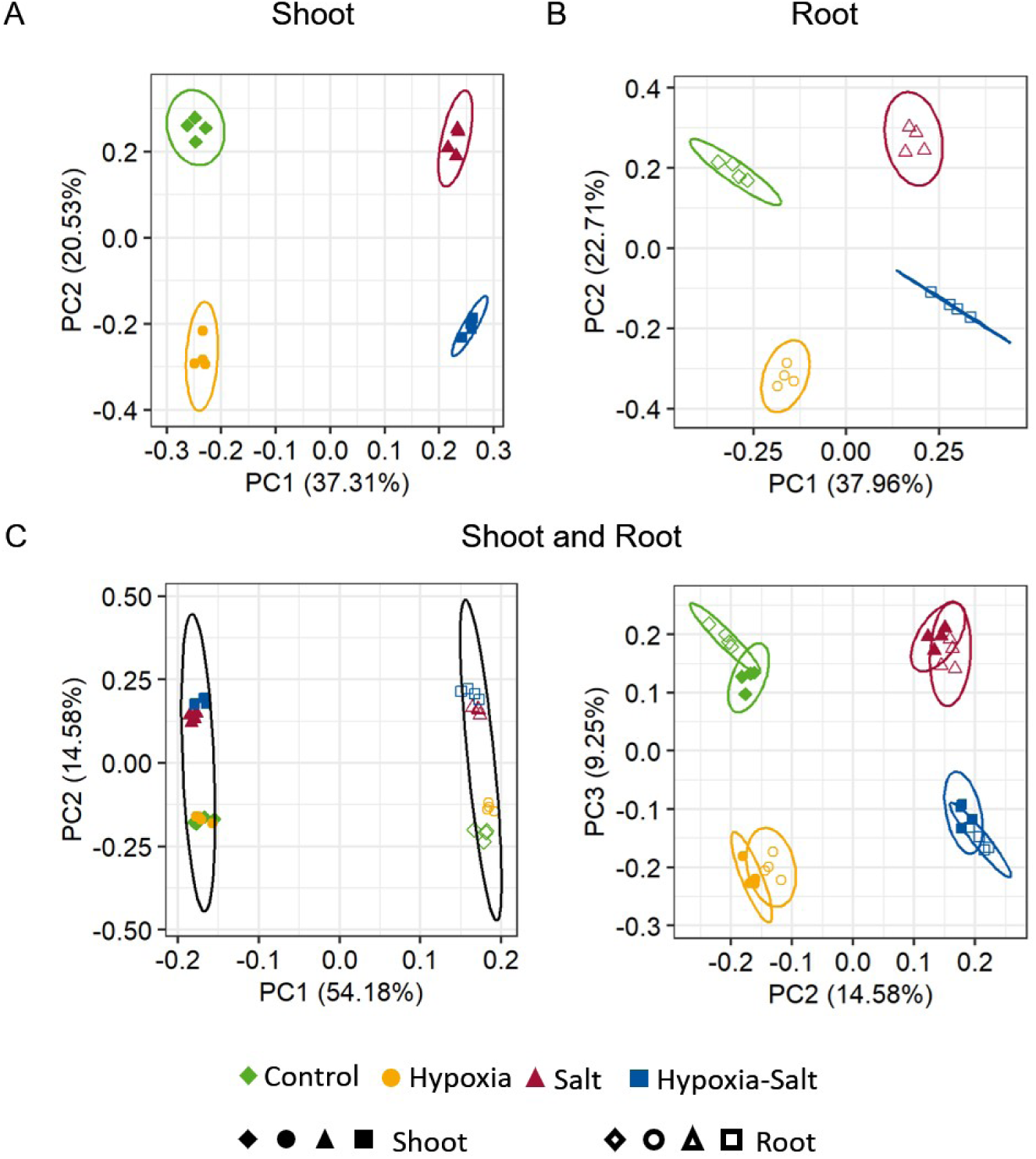
Principle Component Analysis (PCA) of All Annotated Genes Following RNA Sequencing. Counts were transformed using regularized log (rlog) transformation, analyzed separately for **(A)** shoot and **(B)** root samples and **(C)** their combined dataset. Different experimental conditions are depicted by distinct symbols and colors. filled shapes: shoot samples; empty shapes: root samples; green-diamond: control; yellow-cycle: hypoxia; red-rectangle: salt; blue-square: hypoxia salt.

### Hypoxia-Salt Stress Drives Extensive Reprogramming in Various Metabolic Pathways

In order to investigate how combined exposure to hypoxia-salt reprogram gene expression compared to individual stress conditions, differential gene expression analysis was conducted. Genes were classified as significantly differentially expressed (sDEG) relative to control samples if they had an adjusted p-value*<*0.01 (see Material and Methods, sDEGs). Across all conditions, roots exhibited a higher total number of sDEGs compared to shoots (Figure 2A, blue and grey numbers, Suppl. Fig. S6B). Simultaneous hypoxia-salt treatment yielded the highest number of sDEGs, with 3370 genes affected in shoots and 3877 in roots, surpassing other conditions. Subsequently, these substantial numbers indicate extensive gene reprogramming under hypoxia-salt, as approximately half of the genes were significantly changed (in shoots ∼48% and in roots ∼54%). For the individual stresses, salt affected 39-43% (shoot-root), while hypoxia influenced 23-30% (shoot-root) genes (Suppl. Fig. S6B, Fig. 2A, blue and grey numbers). In addition, comparison of up-regulated versus down-regulated transcripts revealed nearly balanced numbers, with slightly more up-regulation under hypoxia-salt and hypoxia treatments (Suppl. Fig. S6B). Subsequently, comparing numbers of highly up- and down-regulated gene expressions (abs(log2FC)*>*2) revealed a higher number of up-regulation in hypoxia-salt and hypoxia compared to down-regulation (Fig. 2A, blue).

**Figure 2:**
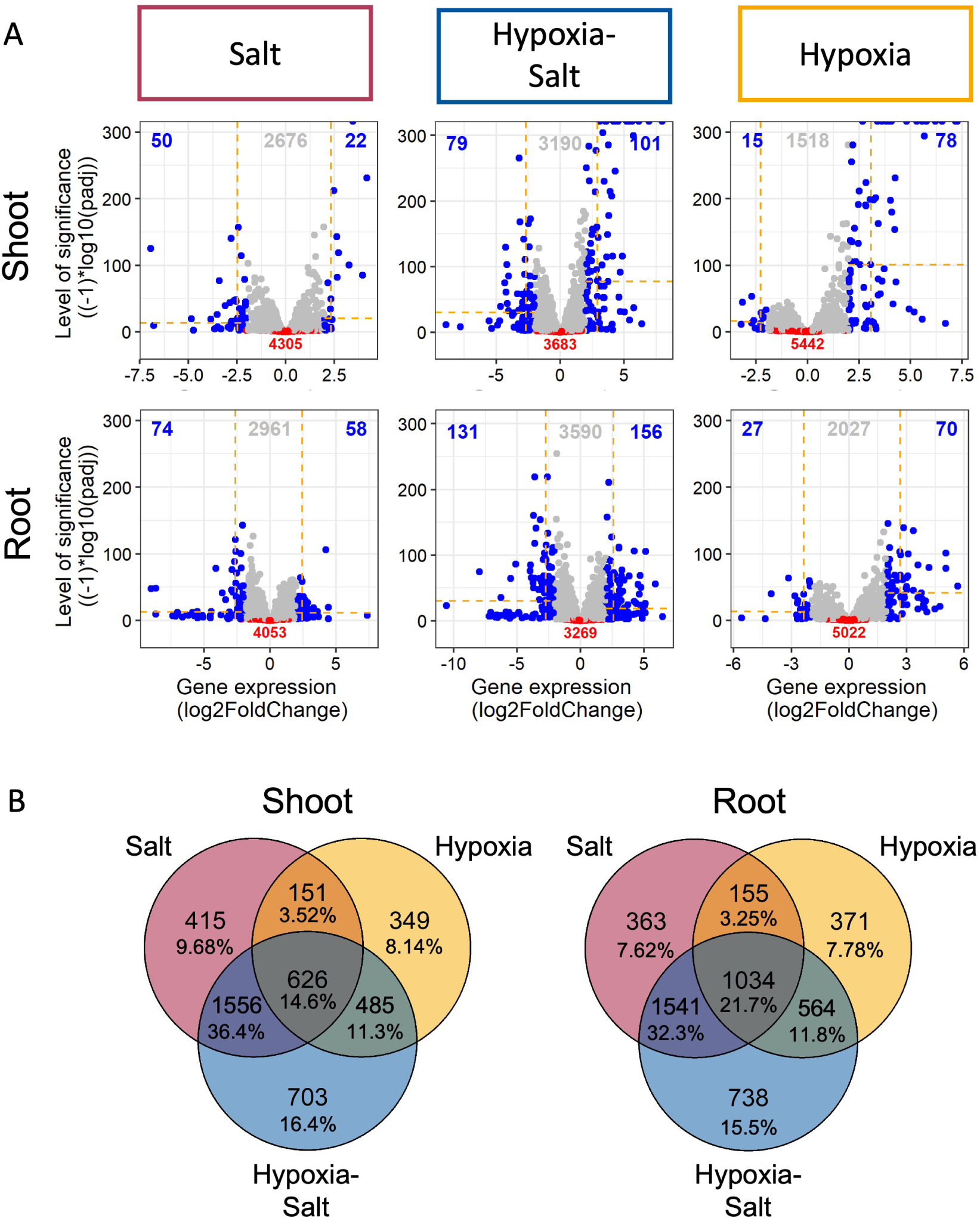
Differential Gene Expression Analysis under Salt, Hypoxia-Salt and Hypoxia in Shoots and Roots. **(A)** The log2FoldChanges (log2FC) of gene expression for shoots and roots were plotted against their significance level (-log10 of the adjusted p-values) across different conditions in both tissues. Blue dots: significant differentially expressed genes (sDEGs) with high log2FC values (p-value*<*0.01 and abs(log2FC)*>*2)). Grey dots: sDEGs with a lowered log2FC (p-value*<*0.01 and abs(log2FC)*<*2). Red dots: genes without significant changes (p-value*>*0.01). Colored numbers indicate the number of genes in their corresponding category. Yellow dashed lines mark the median of the gene expression change (log2FC, vertically) and the median of the significance level ((-1)*log10(padj), horizontally) for highly regulated genes (blue dots). **(B)** The overlap of sDEGs (significant differentially expressed genes, p-value *<*0.01) among individual hypoxia, salt, and simultaneous hypoxia-salt stress. Each circle represents the sDEGs set for one condition, with overlapping areas indicating shared genes between conditions, while unique genes are shown in non-overlapping sections. Abbreviations: abs:= absolute; log2FC:= logarithmic fold change in gene expression; DEGs:= differentially expressed genes; sDEGs:= significant differentially expressed genes; p-value:= significance value; padj:= adjusted p-value.

Genes exhibiting high log2FCs in their expression were analyzed and filtered from the data (Suppl. File 1, Sheet ’Volcano_highly_regulated_DEGs’, Suppl. Fig. S9 and S10). In both tissues during hypoxic conditions (**H**, **HS**), highly up-regulated transcripts included genes like *SUCROSE SYNTHASE (SUS), ETHYLENE RESPONSIVE FACTORS group VII* (*ERFVII; RAP2.12*), *PLANT CYSTEINE OXYDASE (PCO)* and *PYRUVATE DECARBOXYLASE (PDC)* (Suppl. Fig. S9 and S10). During salt treatments (**S** and **HS**), *SUGAR WILL EVENTUALLY BE EXPORTED TRANSPORTER (SWEET)* paralogs exhibited both strong up-regulation and down-regulation across tissues. Under hypoxia alone (**H**) *SWEETs* were not impacted severely in expression changes (Suppl. Fig. S9 and S10). Further carbohydrate metabolism related genes varied in expression based on tissue type and condition. For example, *GLUCOSE-6-PHOSPHATE DEHYDROGENASE (G6PDH)* and *ALDEHYDE DEHYDROGENASE 2 (ALDH2)* were strongly down-regulated in their expression under saline conditions (**S** and **HS**) in shoots.

Under simultaneous stress (**HS**) paralogs of *TREHALOSE-6-PHOSPHATE PHOSPHATASE (TPP)* displayed both high up- and down-regulation in the gene expression of the shoots, and were up-regulated in roots. *VACUOLAR/CELL WALL INVERTASE INHIBITOR (VIF2)* expression was down-regulated in shoots and *FRUCTOSE-1,6-BISPHOSPHATE ALDOLASE (FBA1)* expression was up-regulated in roots under simultaneous stress. Independent of tissue or conditions, the expression of branched-chain amino acid degradation genes were down-regulated, with the exception that no amino acid-related gene was highly regulated in shoots under hypoxic conditions. Genes involved in GABA (gamma-aminobutyric acid) and methionine degradation were down-regulated under simultaneous hypoxia-salt in both tissues, as well as under salinity alone (**S**) in the roots. Interestingly, *GLUTAMATE DECARBOXYLASE (GAD)* (Suppl. Fig. S9 and S10) a gene involved in the biosynthesis of GABA was up-regulated in the shoots under hypoxic conditions (**H** and **HS**), and in the roots under simultaneous hypoxia-salt.

Additionally, in regard of energy metabolism, a hypoxia specific up-regulation of *SUCROSE NON-FERMENTING 1-RELATED PROTEIN KINASE 3* (*SnRK3*) was observed in shoots (**H** and **HS**), and in roots under **H**. Overall, the reprogramming after simultaneous hypoxia-salt treatment revealed the highest significant changes in differential gene expression patterns independent of the examined tissue, indicating a unique effect that enhances the simultaneous response compared to each condition alone (Fig. 2A, grey and blue). To evaluate whether simultaneous stress affects specific functional categories, we conducted an enrichment analysis using MapMan across all metabolic pathways (Schwacke et al., 2019). Enriched categories represent metabolic pathways that are over- or under-represented in experimental conditions compared to a reference group. Key enriched categories (p-value *<* 0.05) in shoots under all conditions included i) cellular respiration, ii) coenzyme metabolism, iii) photosynthesis, iv) protein biosynthesis (Suppl. Fig. S11). Under hypoxia-salt conditions, no unique gene categories were significantly enriched beyond those impacted by individual stresses in both shoots and roots. Notably, secondary metabolism was enriched in roots under single stressors but not combined stress, suggesting potential antagonistic effects.

Next we explored the extent to which the differential gene expression under simultaneous stress conditions reflects the combined effects of individual stresses (Fig. 2B). Overall, ∼15% (626 sDEGs) of shoot genes and ∼22% (1034 sDEGs) of root genes showed significant changes regardless of the conditions type (Fig. 2B, total overlap). As expected due to experimental setup and PCA analysis (Fig. 1) in shoots and roots, salt treatments (**S**, **HS**) shared the highest proportion of sDEGs, with 1556 shoot genes (∼36%) and 1541 root genes (∼32%). The overlap between hypoxia and hypoxia-salt was lower with ∼11% in shoot and ∼12% in root. The lowest proportion of sDEGs was shared between salt and hypoxia with ∼4% from shoots and ∼3% from roots. Unique sDEGs proportions varied across conditions for shoot and root in i) salt ∼8 - 10% ii) hypoxia-salt ∼15 - 16% and iii) hypoxia ∼8% (Fig. 2B). As **HS** was most impacted, there nearly equal proportions of shoot sDEGs were either up- (∼16%) and down-regulated (∼15%) uniquely (Suppl. Fig. S8). In the root **HS**, slightly higher proportions of up-regulated genes (∼17%) than down-regulated sDEGs (∼13%) were found to be unique for hypoxia-salt (Suppl. Fig. S8). Taken together, hypoxia-salt (**HS**) led to the most unique significantly differentially expressed genes, highlighting that the combination of both conditions impacts gene expression distinctly.

### Hypoxia-Salt Responses can be Driven by Synergistic but also Antagonistic Effects of Hypoxia and Salt Responsive Pathways

To investigate whether the combined hypoxia-salt response arises from synergistic or antagonistic interactions, we compared transcriptional changes under single and combined stress conditions.

We analyzed how the summed individual stress responses equal the combined hypoxia-salt changes (log2FC(**H**+**S**)=log2FC(**HS**), log2FoldChanges (log2FC)). Deviations from expected additive outcomes were classified as either synergistic or antagonistic effects. To evaluate whether these effects are predominantly influenced by uniquely hypoxia-salt specific genes or highly expressed genes, we examined their respective proportions.

In the volcano plot analysis of hypoxia-salt related significant differentially expressed genes (sDEGs) (Fig. 3A), over 82% were classified as additive in both shoots and roots. Synergistic and antagonistic effects each represented ∼7% of sDEGs in shoots, and in roots these proportions increased slightly to ∼9%. Interestingly, highly expressed genes showed predominant synergistic or antagonistic effects, collectively accounting for over 85% of sDEGs across both tissues. In contrast, genes uniquely expressed under hypoxia-salt stress (Fig. 3B) exhibited less than 15% involvement in synergistic or antagonistic expression within either tissue type.

**Figure 3:**
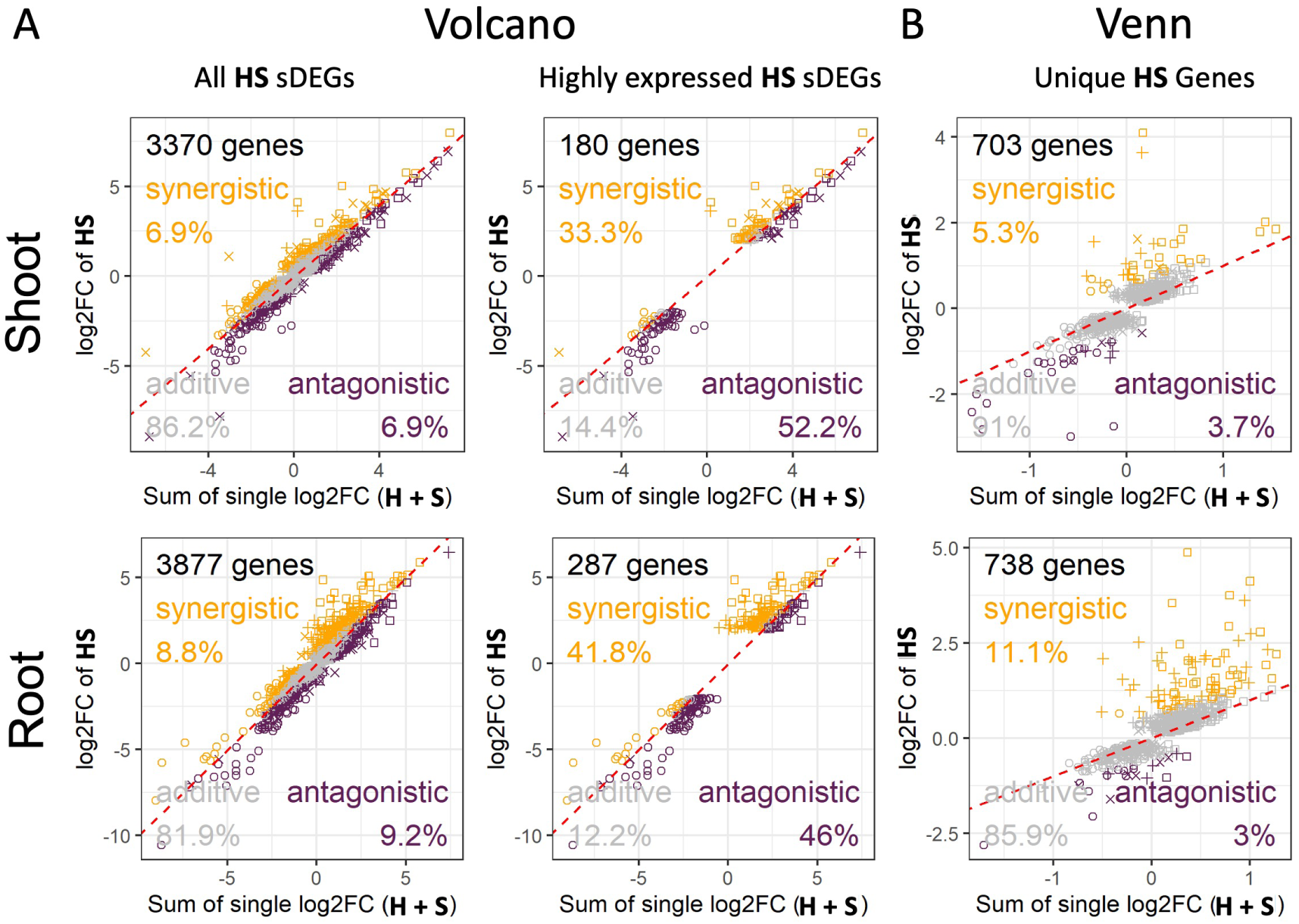
Deviation of Significant Gene Expression Responses from Additive Effects Under Combined Hypoxia-Salt Stress. The relationship between the summed log2FoldChanges (log2FC) of individual stress responses (salt and hypoxia) and the log2FC under simultaneous hypoxia-salt (**HS**) stress in both tissues for **(A)** all significant differentially expressed genes (sDEGs) and highly expressed sDEGs of the hypoxia-salt response (Volcano analysis, Fig. 2A grey and blue dots) as well as for **(B)** uniquely (only in **HS**) sDEGs hypoxia-salt genes (Venn analysis, Fig. 2B). Grey markers denote genes with additive effects, while orange and violet markers indicate genes with synergistic or antagonistic effects, respectively. Additive effects were defined as follows: when the sum of individual stress responses matches the **HS** response (FC(**HS**) = FC(**H**+**S**) ±FC(0.5) confidence interval). Synergistic effects were defined when **HS** expression levels exceed the sum by at least 0.5 FC, and antagonistic effects, when **HS** was below the sum by 0.5 FC threshold. The red diagonal denote the expected trend for additive responses. Symbols represent the sign of log2FC of the individual stress (circle: both (**H** and **S**) negative; square: both (**H** and **S**) positive; +: **H** negative and **S** positive; ×: **H** positive and **S** negative) Abbreviations: FC:= Fold change; DEGs:= Differentially expressed genes

Only a small number (<30) of shoot- and root-specific uniquely sDEGs were classified as highly differentially expressed synergistic or antagonistic genes (Examples in: Suppl. Fig. S12 and S13), Suppl. File 1, Sheet ’Overlap_highVolcano_UniqueHS’), with most exhibiting low expression levels (counts) and high variance across the four replicates. Most key drivers behind the hypoxia-salt response were also identified as sDEGs in one or both individual stress treatments (Suppl. File 1, Sheet ’Overlap_highVolcano_UniqueHS’). These genes participate in pathways responsive to hypoxia or salt, such as amino acid metabolism, cellular respiration, and carbohydrate metabolism.Therefore we selected these metabolic routes to unravel the synergistic and antagonistic impact in depth. The majority of data points (*>* 80%) cluster near the additive response (Fig. 4A, dashed red line). Cellular respiration exhibited the highest additive values in both shoot and roots (*>* 92%), while shoot amino acid metabolism (*>* 84%) and root carbohydrate metabolism (*>*81 %) had the lowest. Genes with antagonistic expression levels were proportionally higher or at least equal compared to those with synergistic effects. In shoots, gene expression patterns separated as follows: i) amino acid metabolism with ∼7% synergistic versus ∼9% antagonistic, ii) carbohydrate metabolism with ∼4% synergistic versus ∼8% antagonistic, and iii) cellular respiration with ∼4% synergistic versus ∼4% antagonistic. In roots, the patterns revealed a greater antagonistic influence compared to shoots. Root gene expression were distributed for i) amino acid metabolism with ∼3% synergistic and ∼10% antagonistic, ii) carbohydrate metabolism with ∼7% synergistic ∼11% antagonistic and iii) cellular respiration with ∼1% synergistic and ∼5% antagonistic. We then examined 56 genes or gene families specifically known to respond to hypoxia (25 genes) or salt stress (31 genes, Liu et al., 2005; Ma et al., 2018; Yang and Guo, 2018, Suppl. File 1, Sheet ’AddEff_HRG_SRG’). From these, shoots exhibited both stronger synergistic and antagonistic responses compared to roots (Fig. 4B). Specifically, ∼16% of hypoxia- or salt-responsive genes showed synergistic effects in shoots, compared to ∼11% in roots. Antagonistic responses were slightly more pronounced in shoots (∼14%) than in roots (∼13%). These findings underscore that the hypoxia-salt response involves complex interactions between stress pathways, resulting in unique transcriptional outputs that defy predictions based solely on individual stress conditions.

**Figure 4:**
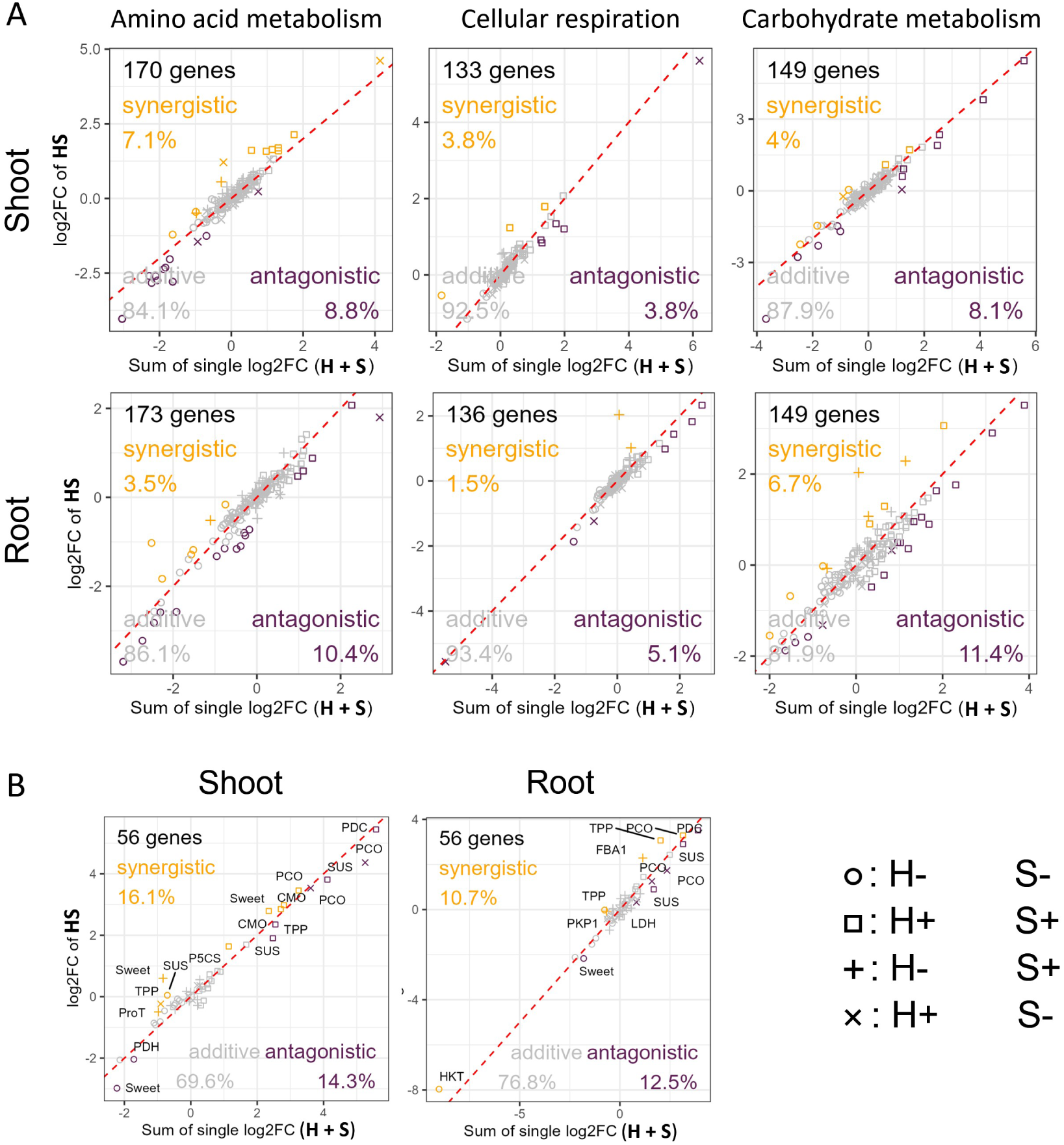
Deviation of Significant Gene Expression Responses from Additive Effects Under Combined Hypoxia-Salt Stress in Hypoxia and Salt Responsive Pathways. The relationship between the summed log2FoldChanges (log2FC) of individual stress responses (salt and hypoxia) and the log2FC under simultaneous hypoxia-salt (**HS**) stress in both tissues for genes of **(A)** amino acid metabolism, carbohydrate metabolism, and cellular respiration pathways as well as for **(B)** known (56 genes) to be involved in hypoxia or salt responses (Liu et al., 2005; Ma et al., 2018; Yang and Guo, 2018, Suppl. File S1 Sheet ’AddEff_HRG_SRG’). Grey markers denote genes with additive effects, while orange and violet markers indicate genes with synergistic or antagonistic effects, respectively. Additive effects were defined as follows: when the sum of individual stress responses matches the **HS** response (FC(**HS**) = FC(**H**+**S**) ±FC(0.5) confidence interval). Synergistic effects were defined when **HS** expression levels exceed the sum by at least 0.5 FC, and antagonistic effects, when **HS** was below the sum by 0.5 FC threshold. The red diagonal denote the expected trend for additive responses. Symbols represent the sign of log2FC of the individual stress (circle: both (**H** and **S**) negative; square: both (**H** and **S**) positive; +: **H** negative and **S** positive; ×: **H** positive and **S** negative) Abbreviations: FC:= Fold change; PCO:= plant cystein oxidase; PDC:= pyruvate decarboxylase complex; TPP:= trehalose phosphatase; SUS:= sucrose synthase; Sweet:= sugars will eventually be exported transporters; CMO:= choline monooxygenase; P5CS:= pyrroline-5-carboxylate synthase; ProT:= proline transporter; LDH:= lactate dehydrogenase; FBA:= fructose-1,6-bisphosphate aldolase; PK:= pyruvate kinase; HKT:= potassium/sodium cation transporter

### Simultaneous Hypoxia-Salt Alters Central Metabolic Pathways

The previously uncovered synergistic and antagonistic responses to hypoxia and salt in shoot primary metabolic pathways were deeper examined. Carbohydrate metabolism analysis (Fig. 5) revealed transcriptional adjustments in sucrose and starch degradation, as well as trehalose metabolism.

**Figure 5:**
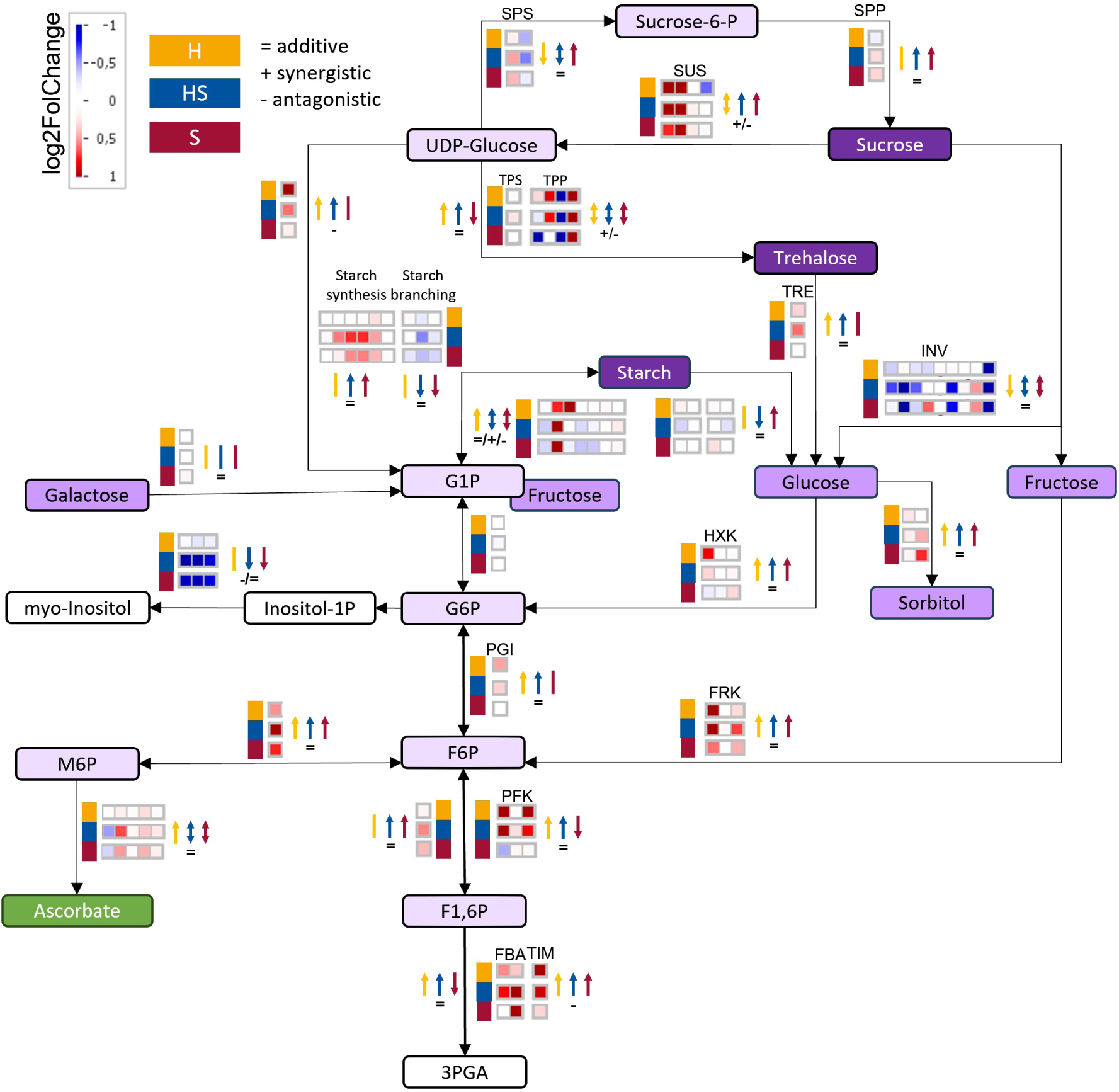
Carbohydrate Pathway Specific Transcriptional Expression Changes Under Hypoxia, Hypoxia-Salt and Salt in *S. europaea*. Expression changes (log2FoldChange, log2FC) are displayed on a custom pathway using the Mapman tool (Schwacke et al., 2019), an assembled reference transcriptome was annotated using Mercator and Mapman as reference. Sugar phosphates, monosaccharides and di-/polysaccharides were marked in light to dark violet, respectively. Organic acids were highlighted in green. Black arrows between the metabolites display the enzymatic conversion. Changes in the transcripts encoding these enzymes are indicated in the boxes next to the linking arrow with positive (red) and negative (blue) log2FC. The squares indicate conditions: yellow-H: hypoxia, blue-**HS**: hypoxia-salt, and red-**S**: salt). Color coded arrows indicate up- or down-regulation of expressions for the color coded respective condition together with indications of additive (=), synergistic (+) and antagonistic (-) **HS** responses. If gene expressions were up- and down-regulated, two-headed arrows were used. Abbreviations: 3PGA:= 3-Phosphoglyceric acid; F6P:= Fructose-6-phosphate; F1,6P:= Fructose-1,6-bisphosphate; FBA:= Fructose-bisphosphate aldolase; FRK:= Fructokinase, G1P:= Glucose-1-phosphate; G6P:= Glucose-6-phosphate; HXK:= Hexokinase, Inositol-1P:= Inositol-1-phosphate; INV:= Invertase; M6P:= Mannose-6-phosphate; PGI:= Phosphoglucose isomerase; PFK:= Phosphofructokinase; SUS:= Sucrose synthase; Sucrose-6-P:= Sucrose-6-phosphate; SPP:= Sucrose phosphate phosphatase, SPS:= Sucrose phosphate synthase; TIM:= Triose-phosphate isomerase; TPS:= Trehalose-6-phosphate synthase, TRE:= Trehalase; TPP:= Trehalose-6-phosphate phosphatase

During hypoxia (**H**) expression of *INVERTASE* (*INV*) was down-regulated, whereas *SUCROSE SYNTHASE* (*SUS*) was up-regulated (Fig. 5, yellow). Salt stress (**S**) exhibited similarly higher *SUS* and a bit more varied *INVs* patterns, and additional expression up-regulation linked to starch synthesis (Fig. 5, red). Reduced expression of starch branching enzymes suggested putative enhanced amylose formation. Combined hypoxia-salt stress (**HS**), led to an intensified down-regulation of the *INVs* sucrose degradation genes, coupled with stronger up-regulation of *SUS*, *TPP* and *TREHALASE* (*TRE*) (Fig. 5, blue). Additionally, three out of four *SUS* were regulated synergistic (1) and antagonistic (2) while the *INV* s showed primarily additive effects (8/9 *INV* genes). Half of the *TPP* genes were synergistic or antagonistic regulated under simultaneous stress (Fig. 5, signs, Suppl. File 1, Sheet ’Genes_Carbohydrate_metabolism’). Additionally, starch synthesis was further enhanced alongside reduced branching enzyme genes once more indicating increased amylose formation. Amino acid (AA) metabolism (Fig. 6, Suppl. File 1, Sheet ’Genes_AS_metabolism’), revealed distinct transcriptional changes across different stress conditions. Salt stress (**S**) triggered induction of the serine-derived AA group, particularly glycine synthesis, whereas hypoxia (**H**) repressed this group by down-regulating cysteine synthesis genes and yet maintained expression for glycine synthesis.

**Figure 6:**
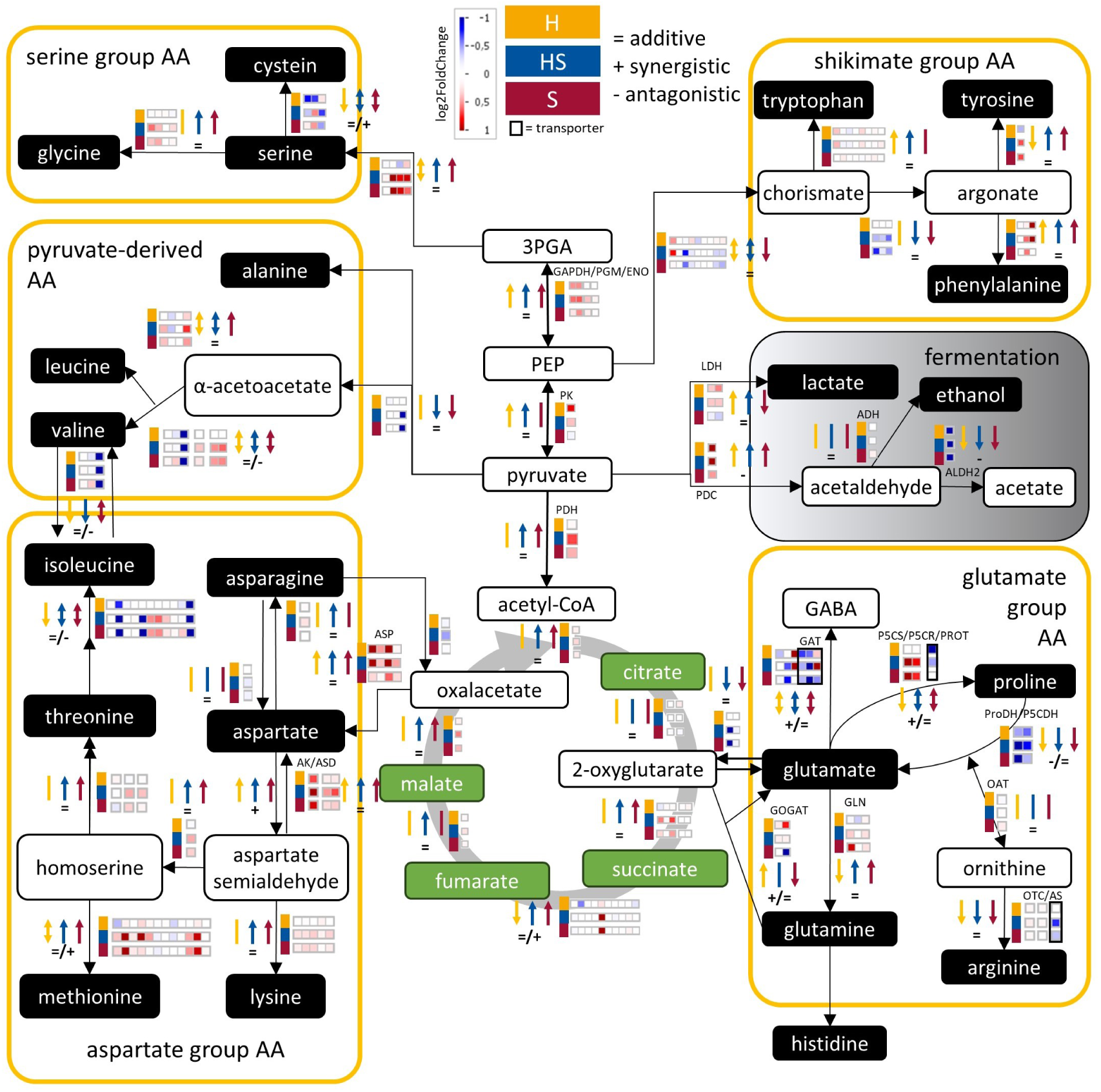
Glycolysis Parts, Tricarboxylic Acid (TCA) Cycle, Fermentation and Amino Acid Metabolism Specific Transcriptional Expression Changes Under Hypoxia, Hypoxia-Salt and Salt in *S. europaea* Shoots. Expression changes (log2FoldChange, log2FC) are displayed on a custom pathway using the Mapman tool, an assembled reference transcriptome was annotated using Mercator and Mapman as reference. Organic acids (green), amino acids and fermentation products (black) and other metabolites (white) are depicted in their metabolic routes. The yellow boxes summarize all amino acids and conversions of the respective amino acid group. Black arrows between the metabolites display the enzymatic conversion. Changes in the transcripts encoding these enzymes are indicated in the boxes next to the linking arrow with positive (red) and negative (blue) log2FC. The squares indicate conditions: yellow-H: hypoxia, blue-**HS**: hypoxia-salt, and red-**S**: salt). Color coded arrows indicate up- or down-regulation of expressions for the color coded respective condition together with indications of additive (=), synergistic (+) and antagonisitc (-) **HS** responses. If genes were up- and down-regulated, two-headed arrows were used. Abbreviations: 3PGA:= 3-phosphoglyceric acid; AA:= Amino acids; acetyl-CoA:= Acetyl-coenzyme A; ADH:= Alcohol dehydrogenase; ALDH2:= Aldehyde dehydrogenase; AK:= Aspartate kinase; AS:= Argininosuccinate synthase; ASD:= Aspartate-semialdehyde dehydrogenase; ASP:= Aspartate aminotransferase; ENO:= Enolase; GABA:= Gamma-aminobutyric acid; GAT:= GABA transporter; GAPDH:= Glyceraldehyde 3-phosphate dehydrogenase; GDH:= Glutamate dehydrogenase; GLN:= Glutamine synthetase; GOGAT:= Glutamate synthase; LDH:= Lactate dehydrogenase; OAT:= ornithine aminotransferase; P5CS:= Pyrroline-5 carboxylate synthetase; P5CR:= Pyrroline-5 carboxylate reductase; P5CDH:= Pyrroline-5 carboxylate dehydrogenase; PDC:= Pyruvate decarboxylase; PDH:= Pyruvate dehydrogenase; PEP:= Phosphoenolpyruvate; PGM:= Phosphoglycerate mutase; PK:= Pyruvate kinase; ProT:= Proline transporter; ProDH:= Proline dehydrogenase

Combined hypoxia-salt (**HS**) serine-derived genes showed additive induction. Shikimate pathway responses varied, salt stress down-regulated early biosynthetic gene expressions (not **H**) but up-regulated phenylalanine, and tyrosine synthesis (Fig. 6, red). Hypoxia up-regulated phenylalanine and tryptophan and down-regulated tyrosine synthesis (Fig. 6, yellow). Combined **HS** caused stronger changes, tryptophan followed the hypoxia **H** pattern (up), tyrosine the salt pattern (up), and phenylalanine showed a mixed response, resulting overall in additive expression across the shikimate group. Pyruvate-derived AA genes were only slightly affected by all treatments, except valine biosynthesis, which seemed to be regulated under combined **HS** stress (Fig. 6, blue).

Gene expressions within the aspartate group AA metabolism - including methionine, threonine, asparagine, and lysine - were up-regulated under both salt (**S**) and simultaneous hypoxia-salt (**HS**) stress. *ASPARTATE AMINOTRANSFERASE* (*AGT*) showed strong induction under hypoxia and additive expression under combined stress, while *ASPARTATE KINASE* (*AK*) was activated by both single stresses and displayed enhanced synergistic effects under simultaneous hypoxia-salt treatment. Salt stress (**S**) induced higher expression patterns in the glutamate group by proline (*P5CS*) and glutamine *via* glutamate, while *GABA TRANSPORTER* (*GAT*), arginine transporter, and *PROLINE TRANSPORTER* (*ProT*) were lowered. This pattern was intensified under simultaneous **HS** stress. Under hypoxia, opposing patterns to salt stress were observed. Genes of the glutamate-derived group responded the most to all treatments. Salt and hypoxia showed contrasting patterns, with salt (**S**) causing higher expressions of *PYRROLINE-5-CARBOXYLATE SYNTHASE* (*P5CS*) and *PYRROLINE-5-CARBOXYLATE REDUCTASE* (*P5CR*), along with lowered expression of the *PROLINE TRANSPORTER* (*ProT*). Proline-degrading enzymes (*ProDH*, *P5CDH*) were repressed under all conditions, whereas the *GABA TRANSPORTER* (*GAT*) was down-regulated by salt (**S**) but induced by hypoxia (**H**). Under combined **HS**, expression patterns resembled those of salt but were more pronounced, with synergistic regulation of *P5CS*, *GAT*, and *ProT*.

## Discussion

Massive reprogramming and adaptation effects of the transcriptome were shown for hypoxia and salt individually for example in *Arabidopsis*, Tomato or Barley Licausi et al., 2011; Liu et al., 2005; Miricescu et al., 2023; Nefissi Ouertani et al., 2021; Roşca et al., 2023; Safavi-Rizi et al., 2020; Yang and Guo, 2017, which are sensitive to both stresses. Yet, the transcriptional reaction to simultaneous hypoxia and salt remains largely unexplored - a gap that limits our understanding of complex stress interactions in natural environments. We wanted to identify gene expression changes in an naturally hypoxia-salt adapted plant by examining synergistic, antagonistic, and additive effects resulting from individual hypoxia or salt treatments. We selected *Salicornia europaea*, known for thriving under fluctuating oxygen and salt levels (Jordine et al., 2024), offering a model for investigating these dynamics. Our comprehensive transcriptome study characterized the gene expression profile shifts of *S. europaea* under hypoxia, salt, and simultaneous hypoxia-salt. Additionally, we aimed to deepen the understanding of the cross-talk between these two stress responses, enabling us to derive first physiological insights into interactions and adaptive mechanisms during simultaneous hypoxia-salt exposure. Although no full genome sequencing data was available for the alignment of the RNA sequencing reads during the time of analysis, a high alignment rate (shoot >63% and root >54%) was achieved with the transcriptome constructed on previously released data (NCBI: SRR822929, SRR823398, SRR944677, SRR944676). Nevertheless, it can not be excluded that some transcripts are not represented in this *de-novo* assembled transcriptome.

### Adapted *Salicornia* Shows Similar Trends as Non-Adapted Plants During Single Stress

Investigations on simultaneous stress has significantly increased over the last years. Gene expression studies have primarily focused on a combination of light, heat, cold, drought, and salinity (Balfagón et al., 2019; Shaar-Moshe et al., 2017; Zheng et al., 2016). However, the impact of combined hypoxia and salinity on gene expression was not thoroughly explored yet. In a controlled hydroponic system, we examined the gene expression response of *S. europaea* to both individual and simultaneous salt and hypoxia stress conditions. Principal component analysis (PCA) clustering (Fig. 1) along with higher number of significant differentially expressed genes (sDEGs) under simultaneous stress (Fig. 2), demonstrates that the combined conditions cause a unique stress response in shoots and roots. The PCA of the four conditions (control, hypoxia, salt, and hypoxia-salt) revealed distinct clustering patterns in both tissues (Fig. 1, A and B). PC1 and PC2 explained 56% of the variance in shoots and 59% in roots. Along PC1, salt-treated samples clustered distinctly from non-salt-treated samples ones (37-38%), whereas PC2 separated plants exposed to hypoxia from those not exposed (21-23%). This suggested that salinity may play a dominant role in shaping gene expression responses under combined hypoxia-salt conditions. The concept of a “dominant stressor” has been previously described (Pandey et al., 2015). Within this framework, the simultaneous stress response is mainly driven by the more severe individual stress. However, this interpretation should be approached with caution, as our experimental setup here, with *Salicornia* plants being acclimated to salt similar to nature to avoid salt-shock responses-still revealed distinct clustering patterns under combined hypoxia-salt stresses. This provides initial evidence supporting uniquely differentiated gene expression resulting from simultaneous stress.

Differential gene expression (DEG) analysis (Fig. 2A) reinforced these findings, revealing not only the highest numbers of sDEG under simultaneous stress but also a high significance ratio (Fig. S6B), indicating a broader transcriptional response. These data are in alignment with other studies on combined stress responses, with an altered expression during the combination compared to single stress application (Sewelam et al., 2014; Shaar-Moshe et al., 2017). Both PCA and DEG analysis emphasize the unique profile of the simultaneous hypoxia-salt stress response. The combined shoot-root PCA analysis (Fig. 1C), revealed a clear separation by tissue type accounting for 54% of total variance. Similar separations between shoots and roots have been reported for *Populus* (Sun et al., 2019) and *Arabidopsis thaliana* (Apelt et al., 2021). This separation highlights the distinct molecular and physiological adaptations of both organs to their specialized developmental roles (Hodge et al., 2009). Differential gene expression analysis confirmed these differences (Fig. 2A), showing that roots consistently exhibited a higher number of DEGs across all treatments, emphasizing their role in environmental stress responses (Vives-Peris et al., 2020). Under salt stress, transcriptional expression in shoots appeared broadly repressed, with more down- than up-regulated sDEGs. In contrast, hypoxia triggered a strong transcriptional activation, inducing a large set of genes (Fig. 2A). This contrasting pattern - transcriptional repression under salt stress and activation under hypoxia - aligns with findings from previous studies on salt- and hypoxia-sensitive plant species (Liu et al., 2005; Skorupa et al., 2019) and is here validated in the adapted *Salicornia* plants.

### Simultaneous Hypoxia-Salt Stress Activates Uniquely Expressed Genes

To further characterize the unique profile of simultaneous hypoxia-salt stress responses, we compared the differential gene expression for all three conditions and quantified the overlaps and distinctions (Fig. 2B). The analysis revealed a clear pattern of overlap and disjoin across individual and simultaneous conditions. A set of commonly regulated genes (∼15% in shoot and ∼22% in roots) was identified, which are general stress responses players. The most substantial overlap of sDEGs occurred between salt (**S**) and simultaneous hypoxia-salt (**HS**) conditions, with ∼36% shoot and ∼32% root genes shared, indicating salt as the putative dominant stressor (Pandey et al., 2015), partially surely due to our experimental setup to avoid salt-shock. Additionally, our analysis also identified single salt- and hypoxia-specific gene subsets (in shoots: ∼10% **S** and ∼8% **H**; in roots: ∼8% **S** and **H**) Fig. 2B). Of particular interest was the number of genes uniquely identified as sDEGs under simultaneous hypoxia-salt (**HS**) stress, comprising ∼16% in both tissues. This subset suggests a unique metabolic answer in the gene expression upon combined hypoxia-salt stress that are not, or only to a minor not significant extent, changed under individual stress conditions.

### Simultaneous Stress is Not Simply Additive

As hypoxia-salt led to individual response in gene expression patters, we further delved into the analysis of additive, synergistic and antagonistic effects. We analyzed various subsets of determined **HS** responsive gene i) all DEGs and ii) all highly expressed DEGs (Fig. 2A) and iii) uniquely **HS** expressed genes (Fig. 2B). Under simultaneous hypoxia-salt conditions, highly expressed genes predominantly exhibited dominantly synergistic and antagonistic (Fig. 3A) in contrast to genes uniquely regulated under hypoxia-salt conditions (Fig. 3B).

Under simultaneous hypoxia-salt conditions, highly expressed genes predominantly exhibited synergistic or antagonistic behavior (Fig. 3A), contrasting with those uniquely regulated by hypoxia-salt alone (Fig. 3B). Most highly expressed hypoxia-salt-responsive genes were also differentially expressed under one or both single-stress scenarios. As a result, only eight were categorized as uniquely **HS** and highly differentially expressed genes (Suppl. File 1, Sheet ’Overlap_highVolcano_UniqueHS’)(Suppl. Fig. S12 and S13), these showed low expression levels (count level) with a high variation across replicates. These results implied that simultaneous hypoxia-salt responses were primarily driven by hypoxia and salt responsive genes and their pathways - highlighting amino acid metabolism alongside cellular respiration and carbohydrate metabolism, known to react in hypoxia and salt (Narsai et al., 2011; Patel et al., 2020), as pivotal interaction nodes within broader regulatory networks impacted herein. Consequently, we analyzed how these categories are affected under simultaneous stress conditions.

The majority of genes (*>*80%) exhibited additive effects across amino acid metabolism, carbohydrate metabolism, cellular respiration (Fig. 4A) as well as all other functional plant categories (Suppl. Fig. shoots S14 and roots S15). Nonetheless, non-additive effects - synergistic and antagonistic - were determined in all functional categories (*<*20%, Fig. 4A). Notably, tissue-specific differences emerged: shoots tended to be stronger synergistic compared to roots, e.g. in amino acid metabolism (∼7% in shoots and ∼4% in roots) and cellular respiration (∼4% in shoots and 2% in roots). This may indicate that above-ground tissues might benefit more from coordinated transcriptional reprogramming. Conversely, antagonistic effects were more pronounced in roots, especially impacting amino acid (∼11%) and carbohydrate metabolism (∼10%). These findings suggest prioritization of reprogramming towards certain metabolic pathways essential to ensure adaptation and survival amidst hypoxic and saline soil conditions.

Cellular respiration, on the other hand, exhibited the lowest variation in both shoots and roots, possibly indicating a more conserved regulatory mechanism operative under simultaneous stress. Interestingly, the investigation of 56 genes and/or gene families with well-known responses to upon hypoxia or salt (Liu et al., 2005; Ma et al., 2018; Yang and Guo, 2018, Suppl. File 1, Sheet ’AddEff_HRG_SRG’), revealed a lowered additive response with under 80% in both tissues (Fig. 4B). This implies simultaneous stress causes a higher proportion synergistic or antagonistic effects (*>*20%), in genes well-known to be involved in either response to hypoxia or salt stress. In shoots, synergism (∼16% versus ∼11%) and antagonism (∼14 % versus ∼13 %) was higher compared to roots. Therefore, shoots may leverage combined hypoxia-salt stress effects to enhance adaptive processes.

Roots showed a greater proportion of antagonistic in comparison to synergistic responses, which may indicate their function as a primary site for salt uptake and sensing osmotic stress (C.-F. Wang et al., 2022). Moreover, a systemic signaling transduction from roots to shoots could account for generally higher non-additive effect in the shoots. For hypoxia such a signaling transduction mechanism was described in *A. thaliana* within the carbohydrate metabolism (Hsu et al., 2011). Notably prominent are genes such as *PLANT CYSTEINE OXYDASE* (*PCO*), *TREHALOSE PHOSPHATASE* (*TPP*), *SUCROSE SYNTHASE* (*SUS*) and *SUGARS WILL EVENTUALLY BE EXPORTED TRANSPORTERS* (*SWEET*) which possess multiple paralogs exhibiting distinct additive reactions-suggesting functional redundancy but also flexibility and adaptability during concurrent stresses. The presence of multiple paralogs enable plants to finely tune metabolic pathways and adapt seamlessly amidst varied environmental stresses and/or simultaneous exposures (*PCO* Weits et al., 2022, *TPP* W. Wang et al., 2020, *SUS* Baud et al., 2004, *SWEET* Gautam et al., 2022). In summary, simultaneous hypoxia-salt stress elicits unique gene expression responses, characterized by significant non-additive effects across multiple pathways, highlighting complex interactions beyond simple additive expectations.

### The Combination of Hypoxia-Salt Leads to Enhanced Distributions of Carbon Equivalents in Shoots

Following our initial findings on the distinct gene expression patterns under simultaneous hypoxia-salt stress, we focused on detailed analysis of carbohydrate metabolism, glycolysis, the TCA cycle, fermentation, and amino acid metabolism due to their crucial roles in facilitating adaptive responses and energy distribution during environmental stress conditions. In the carbohydrate metabolism (Fig. 5), different adaptation mechanisms were determined in response to the different stress conditions. For example, gene expression changes for C-equivalents distribution pointed under saline (**S** and **HS**) toward sucrose and starch biosynthesis, while during hypoxia (**H** and **HS**) trehalose and glucose-1-phosphate (G1P) biosynthesis were enhanced. As a result, the combination in **HS** resulted in all of them being higher expressed (Fig. 5, blue arrows). This stress dependent storage and re-mobilization (Ribeiro et al., 2022) seemed to clearly differ in the gene expression analysis between, salt, hypoxia and its combination in the adapted *Salicornia europaea*. Salty conditions (**S** and **HS**) led to higher gene expressions for starch synthesis in combination with lowered expression of starch branching, pointing towards an increased amylose formation. Gene expressions for sucrose formation during salt conditions seemed to be also slightly enhanced pointing towards a maintained balanced of partitioning of assimilates between starch and sucrose which is known to be dependent on environmental conditions (McCormick and Kruger, 2015). Together with the up-regulation of *SUS* accompanied with the down-regulation of *INV* in all conditions, suggests a redistribution of carbon fluxes to enable more efficient energy supply and the usage of energy reserves under oxygen deficiency and salinity (Bailey-Serres et al., 2012; Lu et al., 2019) and to participate in intrinsic carbohydrate and energy signaling network like for the products and enzymes like HEXOKINASES (Moore et al., 2003), glucose and fructose (Giesbrecht et al., 2025). The trend for enhanced hypoxic gene expression for the formation of trehalose showed, half of the hypoxia-salt *TPP*s were synergistic and antagonistic regulated. The four *TPP* of *S. europaea* displayed distinct expression pattern, and similar expression shifts were observed in wheat (Du et al., 2022), pointing towards either a specific roles of the paralogs or a compartment specif expression of the *TPP*s. A compartment specific expression was described for *TPP* (Kerbler et al., 2023), but for *S. europaea TPP* no compartment-specificity was described yet, therefore observed reactions could rely on both compartment- and stress-specific reactions. Nevertheless, our data suggests an increased role of trehalose as a protective molecule against oxidative and osmotic damage (Sarkar and Sadhukhan, 2022) also during hypoxic and salt-hypoxic conditions in *Salicornia*. Taken together, our gene expression data for carbohydrates suggests a strong stress dependent re-mobilization and biosynthesis of storage metabolites as well as production of signaling and protective metabolites specific for each individual stress and their combination in the evolutionary adapted *Salicornia europaea*.

### Hypoxia Lead to Lactate Dehydrogenase Expression Changes but not Alcohol Dehydrogenase in Shoots

Amino acid metabolism, along with connected primary pathways, also exhibited distinct expression patterns under different stress conditions (Fig. 6). Contrasting regulatory patterns were observed in the end of glycolysis (Fig. 6), indicating a condition dependent regulation. Under salt stress (**S**), the gene expression of *PK* (*PYRUVATE KINASE*) was not changed, while under hypoxia (**H**) the expression was enhanced and slightly diminished in hypoxia-salt hinting towards a salt-dependent overwriting of the hypoxic induction when both stresses co-occur. Protein PK is known to be an important mediator in the energy producing step under anaerobic respiration (Podestá and Plaxton, 1991; van Dongen et al., 2011). This could indicate diminished responsive in hypoxia-salt conditions upon the presence of salt and a slightly more enhanced role of *PDH* (*PYRUVATE DEHYDROGENASE*) which was highest expressed in **HS** and slightly more in **S**. Gene expression patterns of the following TCA cycle were mainly unaffected under all of the tested conditions. Fermentative genes were up-regulated *LDH* (*LACTATE DEHYDROGENASE*) and *PDC* (*PYRUVATE DECARBOXYLASE*) under all conditions (higher in **H** and **HS**). The lactate fermentation pathway, mediated by *LDH*, showed a more stress-specific pattern, with activation under hypoxia but repression under salt stress. In contrast, *ADH* (*ALCOHOL DEHYDROGENASE*), which encodes the enzyme responsible for the energy-yielding step of the fermentation pathway, showed no change in expression under any condition. *ADH* is typically considered a hypoxia-responsive gene in other species (Licausi et al., 2010), an induction under low-oxygen conditions would have been expected. This might be an evolutionary adaptation of *Salicornia* to have a higher basic expression or no need at all for a changed gene expression of *ADH* which needs further to be tested. *LDH*, which is recycling the NADH produced by glycolysis in the absence of oxygen (Sweetlove et al., 2000), and its product lactate could be important for *Salicornia* shoots, as similar mechanisms are described for water-logged plants where leaves detoxify root lactate via *LDH* (António et al., 2015). Supplementary, *LDH* is known to increase upon hypoxic condition in roots themselves like e.g. barley and other cereals (Posso et al., 2025) as well as in potato tubers (Sweetlove et al., 2000). Nevertheless, it was shown that the isoforms of *LDH* are crucial for the behaviour upon hypoxic conditions and the output of *LDH* activitites upon hpyoxia are varied between them (Sweetlove et al., 2000). In conclusion, there is evidence of our gene expression analysis, that *Salicornia* shoots react differently from non-adapted plants upon hypoxia with not activating *ADH* but putatively more *LDH*. As a summary, the expression pattern of all fermentative genes under combined stress closely resembled that observed under hypoxia alone, suggesting that oxygen limitation remains the primary driver of anaerobic metabolism during simultaneous hypoxia-salt stress. Rather suspected, in this pathway, the dominant stressor is hypoxia, indicating that the general assumption about one stressor being dominant (Pandey et al., 2015) as well as our analysis (Fig. 2) pointing towards salt, may be misleading, as dominance likely varies between metabolic pathways.

### Saline Conditions Profoundly Influence Amino Acid Gene Expressions

The genes responsible for the biosynthesis of specific amino acids - such as proline, serine, glycine, valine, lysine, leucine, and isoleucine - which are known to accumulate in spinach during salt stress (Di Martino et al., 2003), exhibited increased expression under salty conditions in our study. This suggests under **HS** the regulation via the stressor salt, as some of them were equal or even lower expressed under hypoxia alone. Aspartate synthesis genes (*ASP ASPARTATE AMINOTRANSFERASE*) were up-regulated in an additive manner under simultaneous stress, with being higher under single hypoxia compared to single salt conditions. Interestingly, genes for the derived amino acids were higher under salty conditions. Aspartate derived amino acids (like isoleucine, lysine) play key roles in regulatory signaling pathways and nitrogen assimilation (de la Torre et al., 2013). Their role may be central to coordinating the plants adaptive response under combined hypoxia-salt stress.

Additionally, proline metabolism gene expression was strongly up-regulated under **S** and simultaneous **HS** stress, including synthesis genes (*PYRROLINE-5-CARBOXYLATE SYNTHETASE P5CS* and *PYRROLINE-5-CARBOXYLATE REDUCTASE P5CR*) and in parallel reduced of the proline degradation genes (*PROLINE DEHYDROGENASE ProDH* and *PYRROLINE-5 CARBOXYLATE DEHYDROGENASE P5CDH*). Proline synthesis gene (*P5CS*) also had a synergistic effect under simultaneous stress (Fig. 4B), likewise also *ProT* (*PROLINE TRANSPORTER*) and *GAT* (*GABA TRANSPORTER*) were synergistic regulated under simultaneous stress. Hence, an enhanced proline accumulation under simultaneous stress is likely. Osmoprotection and reduction of oxidative damage would be assumed, as proline functions as an osmolyte and a reactive oxygen species (ROS) scavenger (Ejaz et al., 2020). As a summary, single hypoxia gene expression responses were not enhanced for proline, but enhanced specifically for GOGAT (*GLUTAMATE SYNTHASE*) and one *GABA transporter GAT*. The enhanced *GAT* -Transporter for GABA could be interpreted as activation of the GABA shunt known to be active under hypoxia (van Dongen et al., 2003).

In summary, salt treatment determined the expression levels also under combined hypoxia-salt conditions.

## Conclusion

The investigation of *Salicornia europaea* under simultaneous hypoxia-salt stress reveals unique transcriptional responses shaped by complex interactions between metabolic pathways. Notably, 16% of differentially expressed genes are uniquely altered under combined stress conditions, underscoring the distinct metabolic adaptations involved. The detailed analysis revealed that synergistic and antagonistic interactions significantly impact gene expression changes across key metabolic pathways such as amino acid metabolism, carbohydrate metabolism, and cellular respiration. Although additive effects were predominant, non-additive interactions were particularly pronounced in pathways known for their roles in hypoxia or salt responses. Salt stress emerged as a dominant factor influencing amino acid metabolism, particularly enhancing proline synthesis for osmoprotection. Conversely, hypoxia primarily drives fermentative responses through anaerobic pathways. The combination of these stresses lead to altered carbohydrate metabolism and significant non-additive effects, highlighting adaptive mechanisms for energy distribution and resilience in *Salicornia europaea*. These findings deepen our understanding of plant tolerance to multiple abiotic stresses and provide valuable insights into the molecular basis of adaptation in this species and beyond. Future research should focus on elucidating the specific roles of uniquely differentially expressed genes and their associated proteins and metabolites under simultaneous stress conditions to further unravel these adaptive mechanisms. Additionally, exploring similar responses in other naturally tolerant species could expand our understanding of plant resilience against environmental challenges across diverse ecosystems.

## Acknowledgements

We want to thank members of the Fürtauer and van Dongen groups and especially Martin Siedt for his valuable input into this project and Brigitta Ehrt for technical support. Additionally we want to thank Stephan Kusch (FZ Jülich) and Jiangzhao Qian (Panstruga and Heitkam groups) for sharing their knowledge.

## Declarations

### Author contribution

A.J., J.T.v.D. and L.F. designed the experiments, evaluated the data. A.J. analyzed the data. A.J. and L.F. wrote the manuscript, J.T.v.D. revised the manuscript. A.J. and J.A. performed experiments.

### Data Availability Statement

The raw RNA-Seq data sets produced in this study can be accessed through NCBI under Bioproject ID PRJNA1256208 for shoot data and PRJNA1256210 for root data.

---

**File 1:**
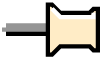
RNA Sequencing Results in .xlsx file format. The file includes comprehensive RNAseq analysis data across sheets ’Shoot_Data’ and “Root_Data”, featuring calculated log2FoldChanges, baseMeans, lfcSE, p-value, padj for each gene under various conditions, along with annotations and additive affiliations. The sheet ’Volcano_highly_regulated_DEGS’ lists genes identified as highly differentially expressed in the volcano analysis. The sheet ’Venn_Unique_Genes’ summarizes genes uniquely associated with one of the stress conditions (**H**, **HS**, **S**) based on Venn analysis. Sheet ’Overlap_highVolcano_UniqueHS’ compiles overlapping genes identified as highly expressed in both volcano plot analyses and uniquely expressed under simultaneous **HS** stress per Venn analysis. The sheet ’AddEff_HRG_SRG’ details hypoxia- and salt-responsive genes utilized in additive effect analyses. All pathway analysis-related genes are compiled within the sheets titled ’Genes_Carbohydrate_Metabolism’ and ’Genes_AA_Metabolism.’

**Figure S1:**
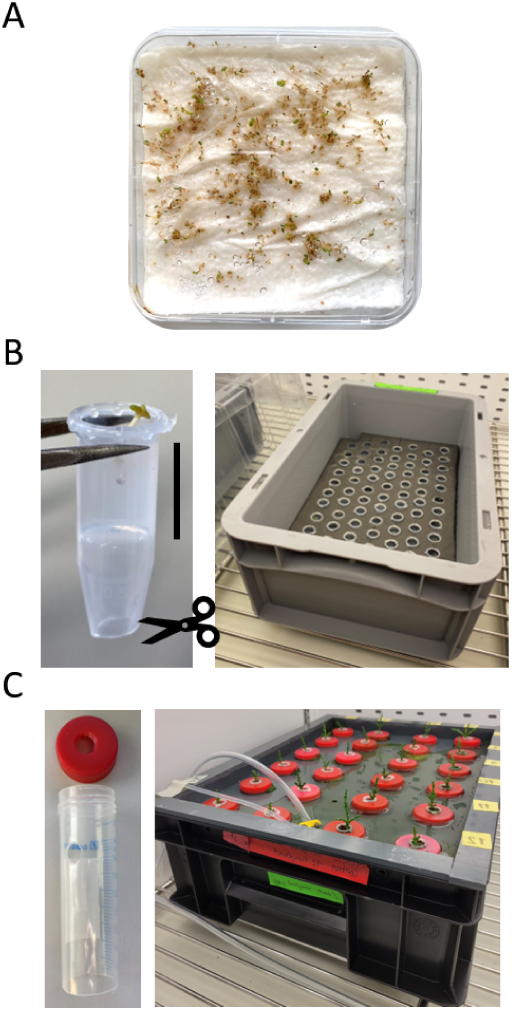
Hydroponic cultivation of *Salicornia europaea.* (**A**) Seeds of *Salicornia europaea* germinated on wet filter paper within a closed square plate for a duration of two weeks. (**B**) Seedlings were transferred to cultivate in liquid 1/2 Hoagland medium during a pre-culture phase lasting three weeks. The upper half (indicated by black bar) of each 1.5 ml reaction tube was filled with solid 1/2 Hoagland medium; tips and lids were removed for planting, allowing insertion into the solid medium for three weeks before further transfer to subsequent tubes. (**C**) Reaction tubes (50 ml) underwent preparation involving tip removal and side holes added to facilitate liquid 1/2 Hoagland medium circulation within the hydroponic system; each plant, along with its initial 1.5 ml reaction tube, was placed within a perforated lid atop these larger tubes. Plants were acclimated for one week before treatment application commenced at the six-week-old stage..

**Figure S2:**
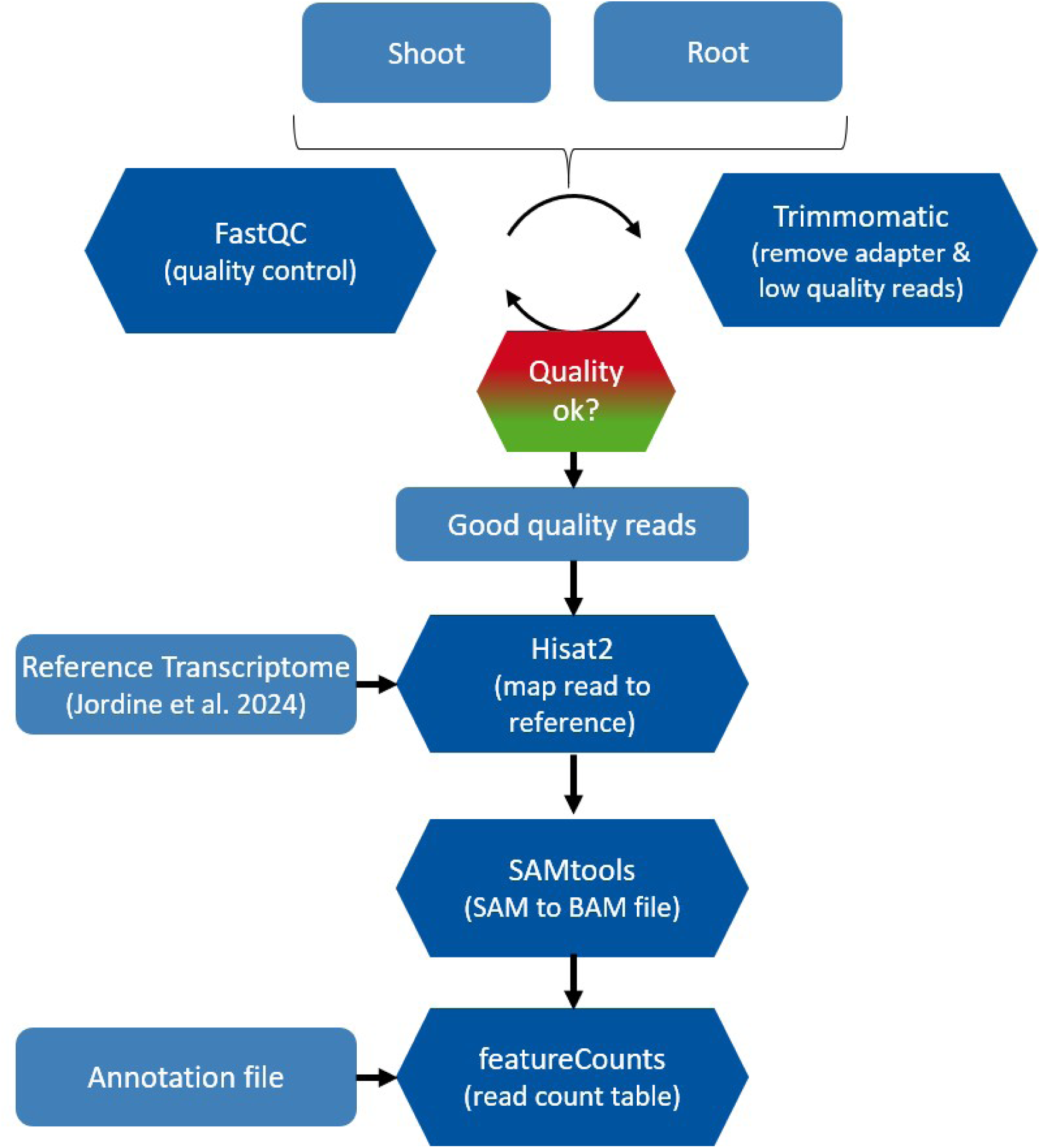
Bioinformatics Workflow for the *in-silico* Processing of Raw Sequences in MiniConda. Quality assessment of raw reads was conducted using FastQC, followed by cleaning to remove low-quality reads and adapter sequences via Trimmomatic. High-quality reads were aligned to a reference transcriptome (Jordine et al., 2024) utilizing Hisat2. Aligned reads were converted from SAM to BAM files, after which a read count table was compiled with annotations using featureCounts.

**Figure S3:**
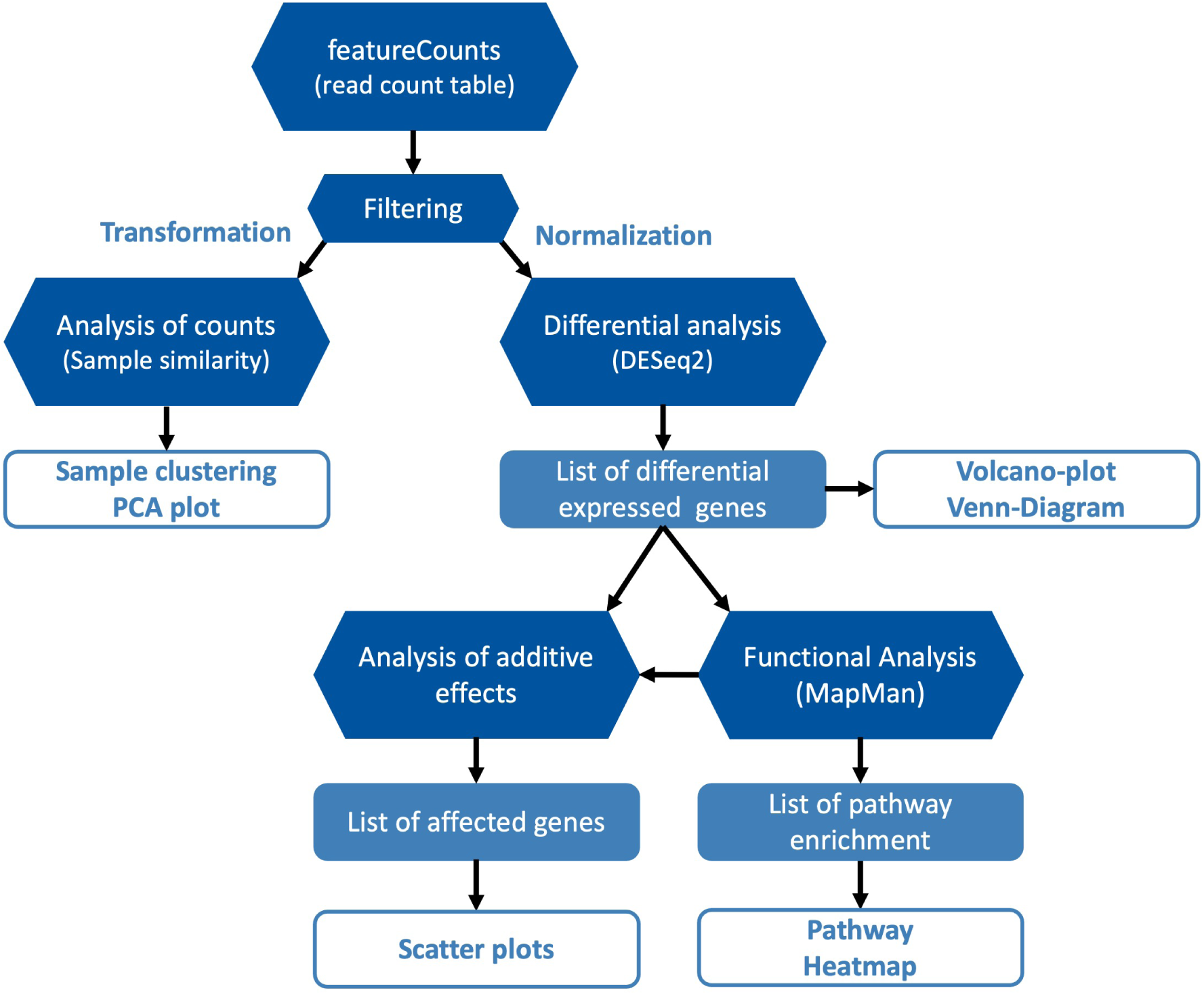
Bioinformatics Workflow of the *in-silico* RNA Sequencing Analysis Conducted in R (https://www.r-project.org/). The read count table was filtered to include counts present in at least three out of four replicate samples. Sample similarity analysis utilized rlog-transformed counts, which were subsequently applied to PCA analysis. Differential gene expression analysis involved normalizing counts using median of ratios normalization, followed by pairwise comparisons via DESeq2, yielding a list of differentially expressed genes (DEGs). Genes with an corrected p-value < 0.01 were classified as significant DEGs (sDEGs). The sDEG list underwent functional analysis using MapMan, producing a categorized gene list. Additive effect analysis was performed on annotated sDEGs within functional categories. Abbreviations: rlog:= regularized-logarithm transformation, PCA:= Principle component analysis, DEGs:= differentially expressed genes

**Figure S4:**
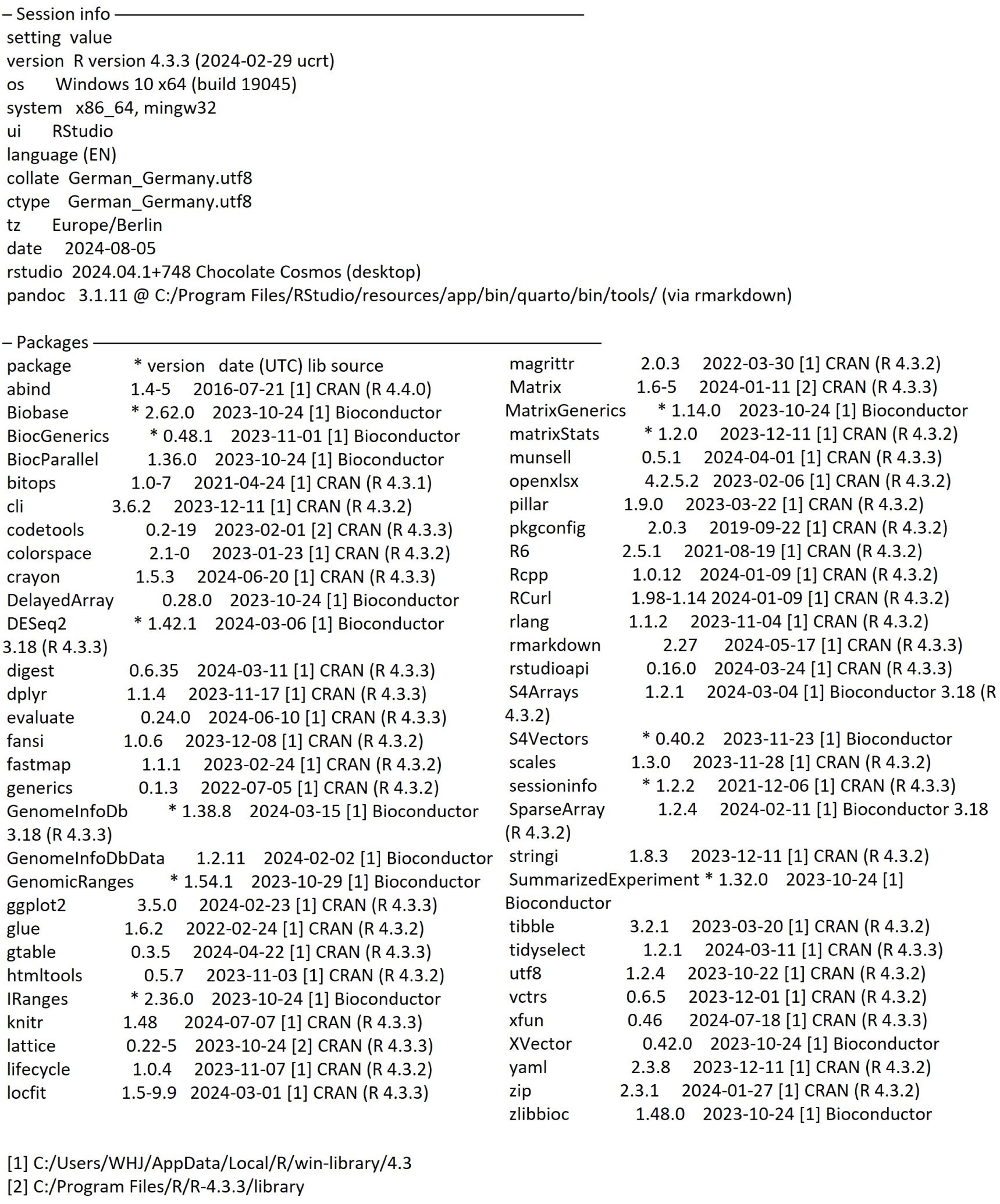
Session Information from RStudio. This session information from RStudio (R Core Team, 2017) details the computational environment utilized for data processing and analysis. The output provides specifics on the R version, operating system, and all loaded packages along with their respective versions.

**Figure S5:**
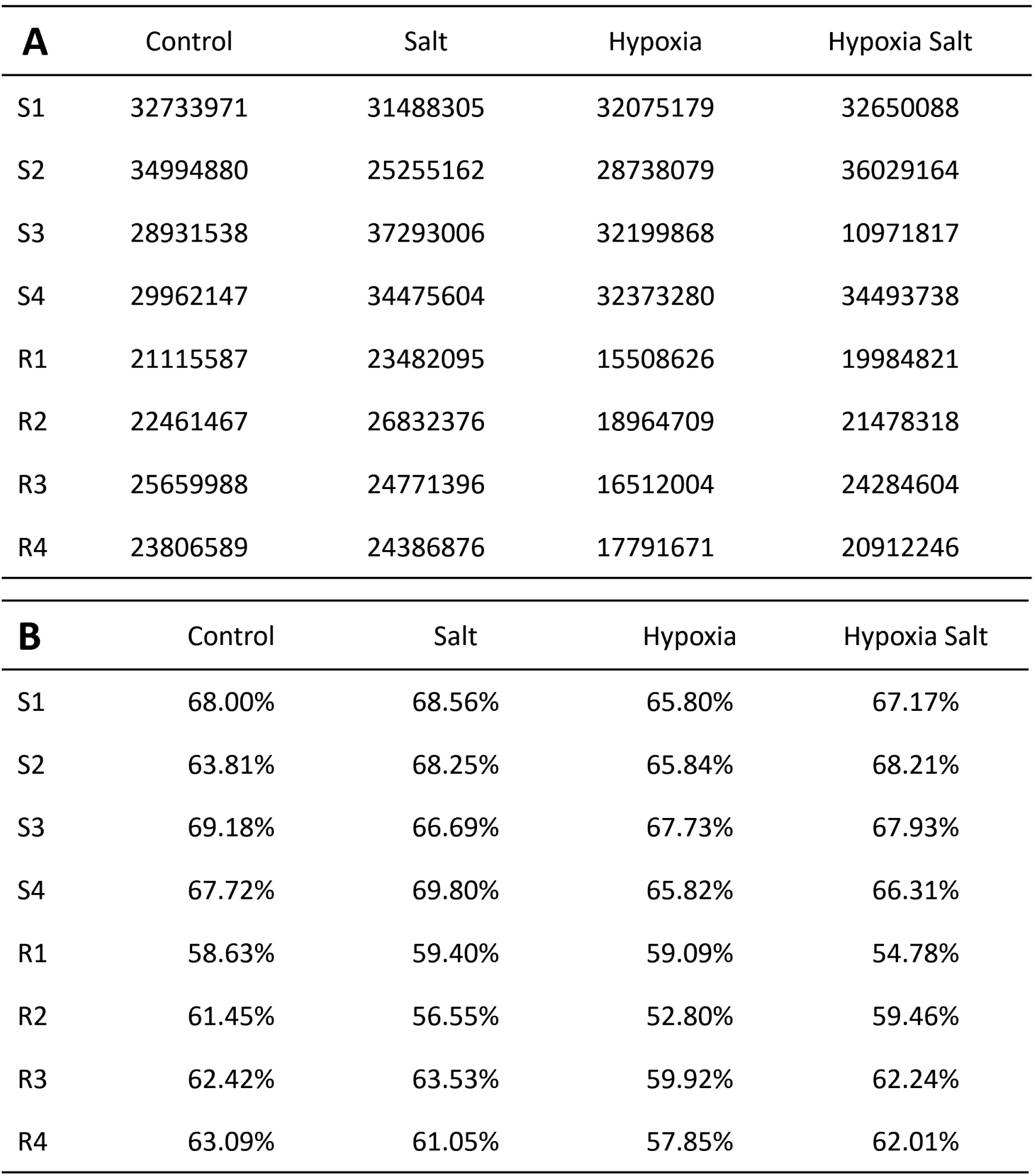
Alignment Information from Hisat2. Hisat2 (Kim et al., 2019) facilitated the alignment of RNAseq reads to the reference transcriptome (Jordine et al., 2024). (**A**) Displays the number of reads per sample, and (**B**) shows overall alignment rates for shoot (S1-S4) and root (R1-R4) samples across different conditions (control: **C**; salt: **S**; hypoxia: **H**, hypoxia-salt: **HS**). Abbreviations: S:= shoot, R:= root, 1-4:= replicate number

**Figure S6:**
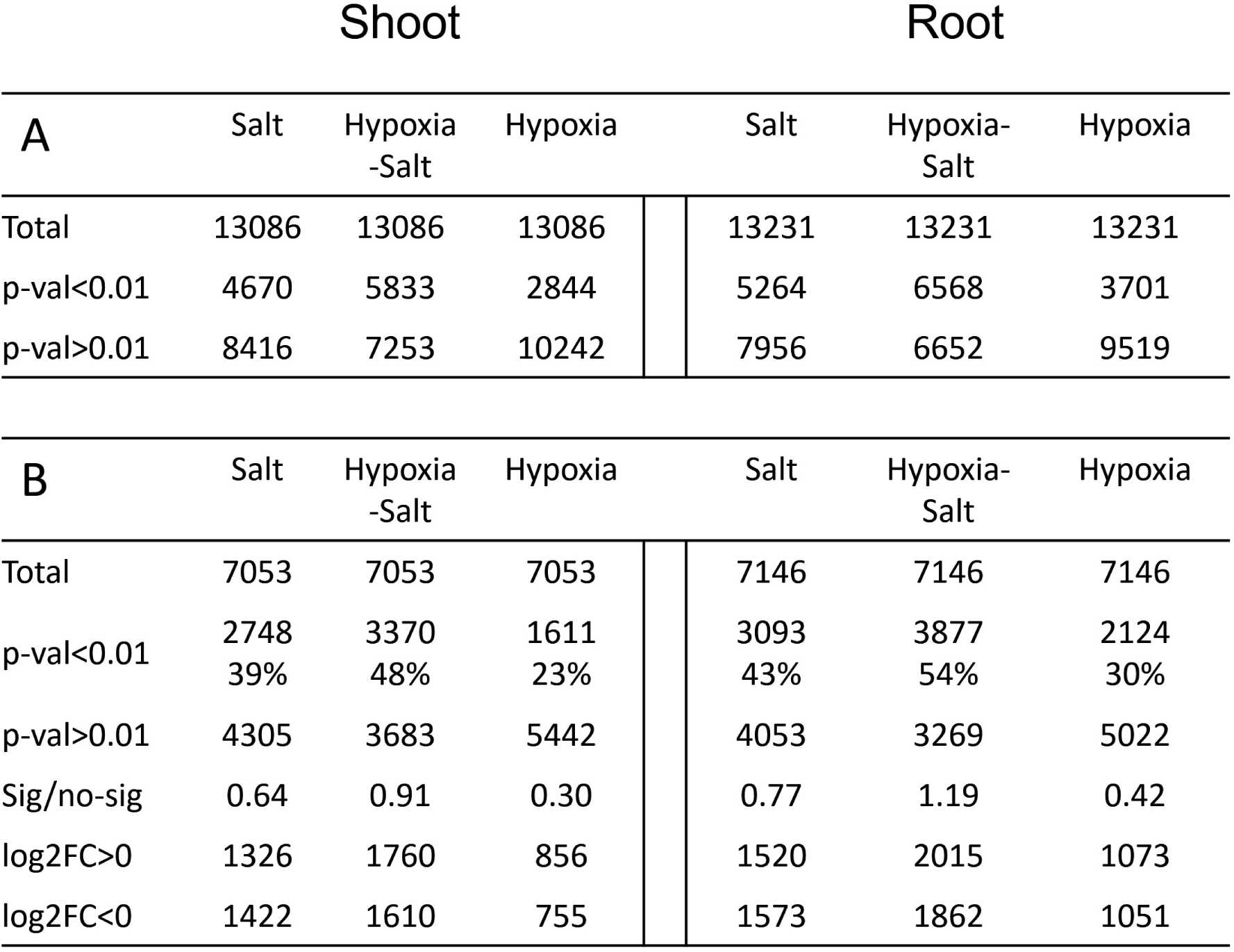
Figures from Differentially Gene Expression Analysis. (**A**) Displays the number of differentially expressed genes, and (**B**) shows differentially expressed annotated genes. ’Total’ represents the complete count of differentially expressed genes. The row p-val*<*0.01 indicates all significant differentially expressed genes (sDEGs) in both numbers and percentages, whereas p-val *>* 0.01 denotes non-significant DEGs. Log2FC *>* 0 captures up-regulated sDEGs, while Log2FC *<* 0 encompasses down-regulated sDEGs. Abbreviations: sDEGs:= significant differentially expressed genes, Log2FC:= logarithmic fold change, p-val:= significance value after correction, sig:= significant, no-sig:= not significant

**Figure S7:**
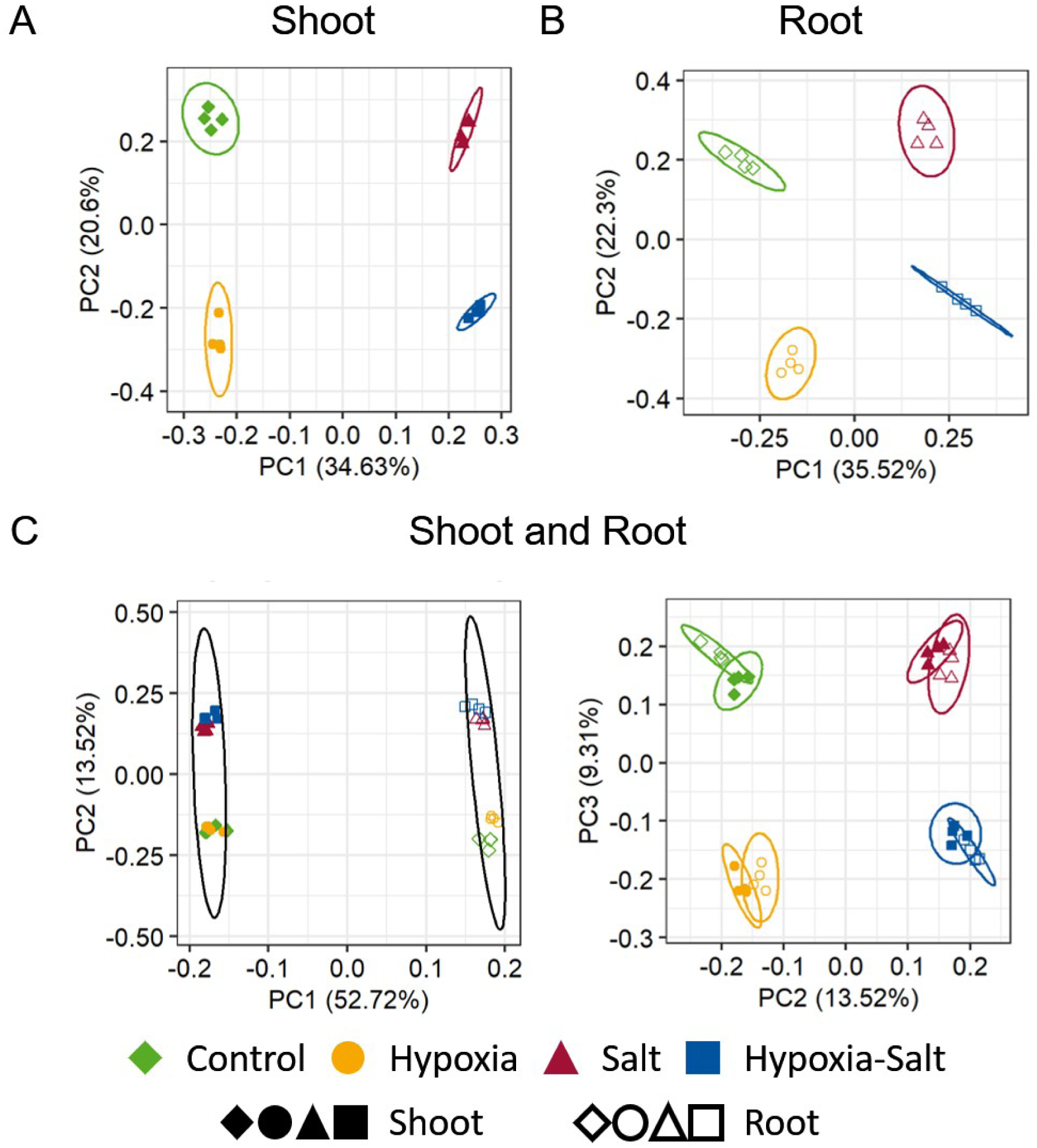
Principle Component Analysis (PCA) of All Genes from RNA Sequencing Datasets. Counts underwent regularized log (rlog) transformation before being analyzed separately for (**A**) shoot samples, (**B**) root samples, and (**C**) the combined dataset. Different experimental conditions are depicted by distinct symbols and colors. filled shapes: shoot samples; empty shapes: root samples; green-diamond: control, yellow-cycle: hypoxia; red-rectangle: salt; blue-square: hypoxia salt,

**Figure S8:**
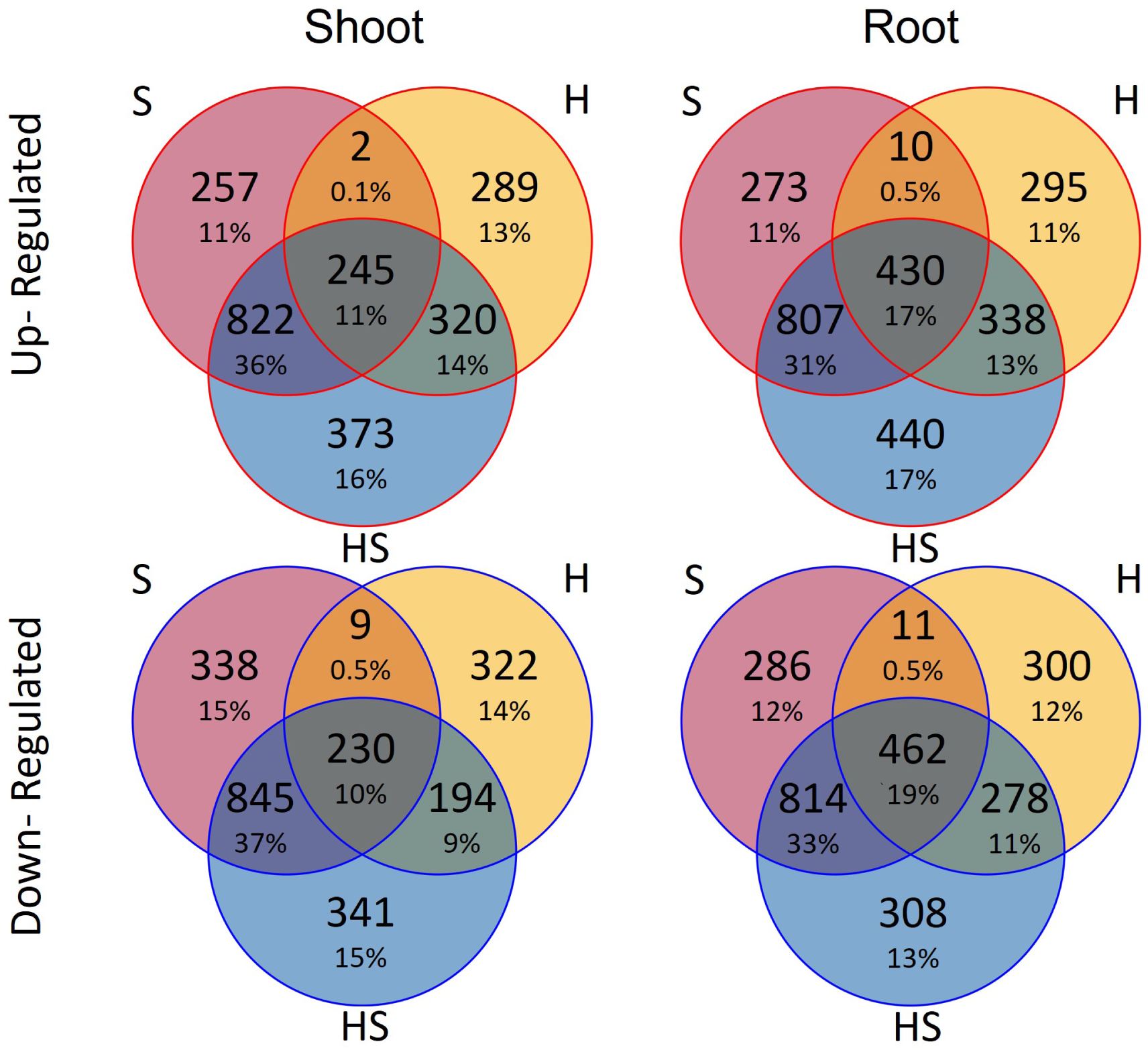
Specificity of the Up- and Down-Regulated sDEGs for the Given Conditions and Their Overlaps. The overlap of sDEGs (significant differentially expressed genes, p-value *<*0.01) across individual hypoxia, salt, and simultaneous hypoxia-salt stress for up-regulated (red border) and down-regulated (blue border) sDEGs. Each circle represents the sDEGs set for one condition, with overlapping areas indicating shared genes between conditions, while unique genes are shown in non-overlapping sections. **H**: Hypoxia (yellow); **HS**: Hypoxia-salt (blue); **S**: Salt (red)

**Figure S9:**
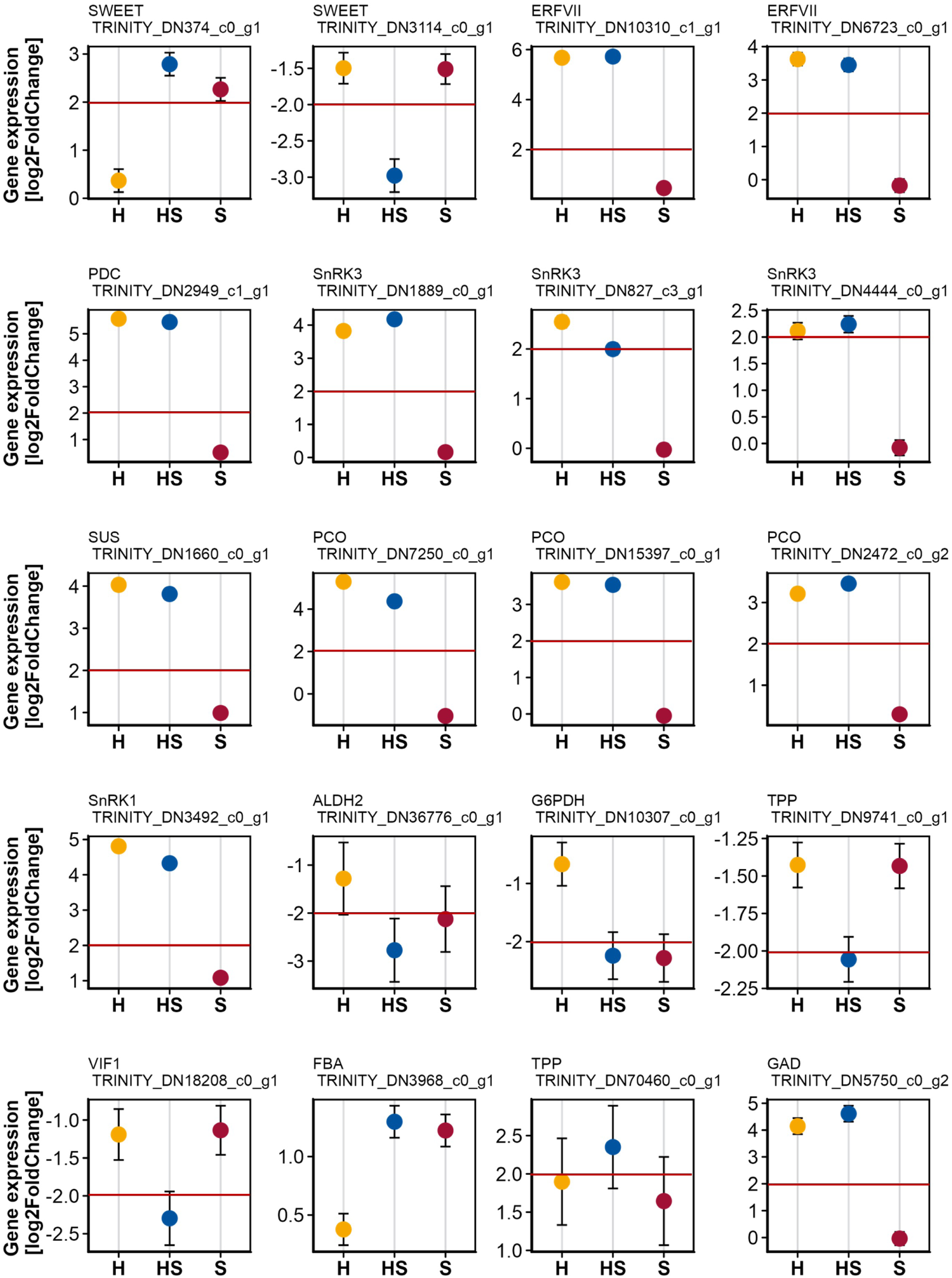
Shoot-Specific Differential Gene Expression of Selected Highly sDEGs. Example genes identified as highly differentially expressed in shoots and/or roots under hypoxia-salt (**HS**) stress (log2FoldChange). Red line marks the log2FC threshold of log2FC*>*2 and log2FC*<*-2. Gene names along with their corresponding TrinityIDs are provided above each graph. **H**: Hypoxia (yellow); **HS**: Hypoxia-salt (blue); **S**: Salt (red) Abbreviations: SWEET:= Sugar will eventually be exported transporter; SUS:= Sucrose synthase; PCO:= Plant cysteine oxidase; ERFVII:= Ethylene responsive factors group VII; PDC:= Pyruvate decarboxylase; ALDH2:= Aldehyde dehydrogenase; TPP:= Trehalose-6-phosphate phosphatase; VIF2:= Vacuolar/cell wall invertase inhibitor; FBA1:= Fructose-1,6-bisphosphate aldolase 1; G6PDH:= Glucose-6-phosphate dehydrogenase; GAD:= Glutamate decarboxylase; SnRK:= Sucrose non-fermenting 1 related protein kinase

**Figure S10:**
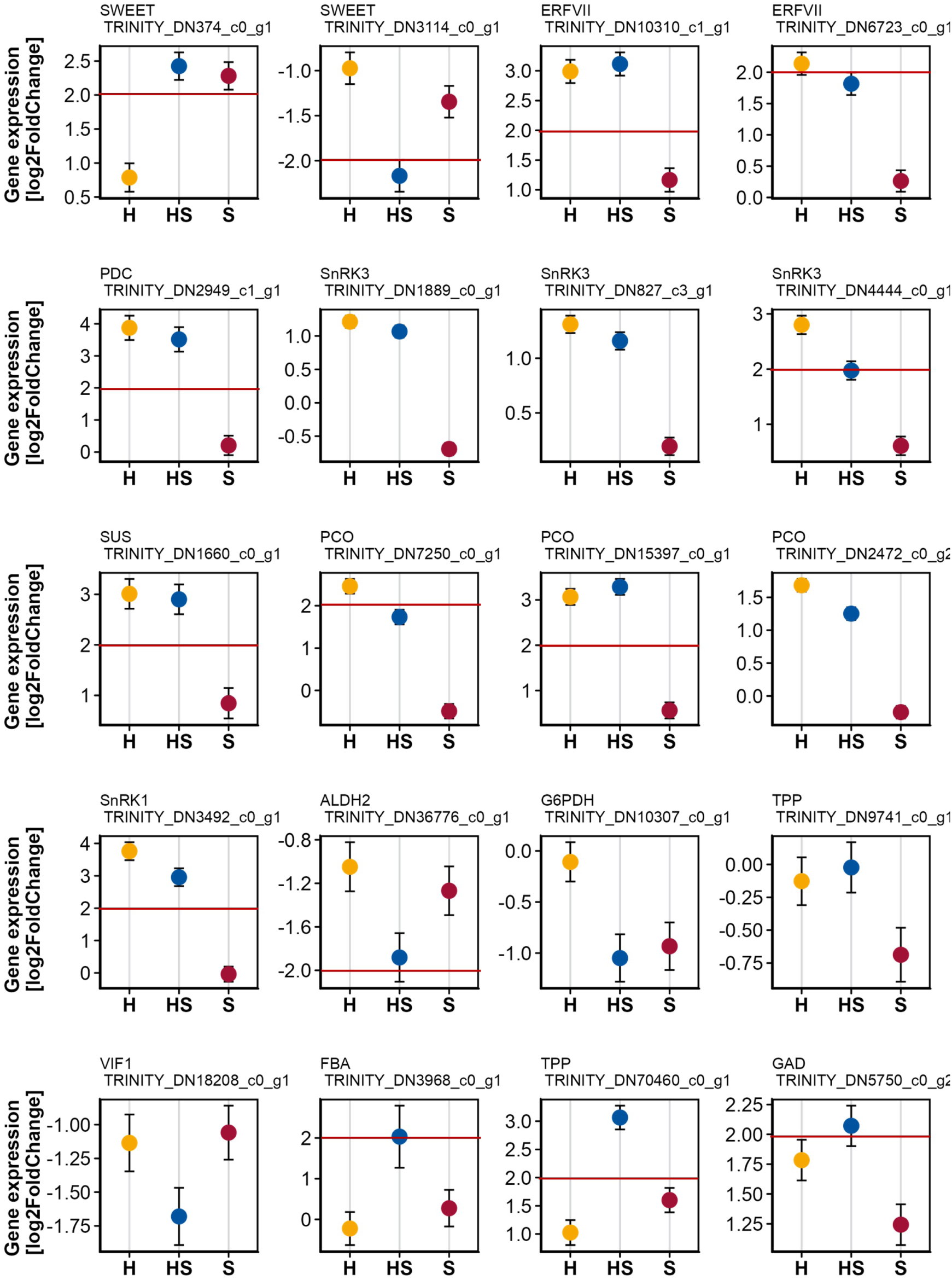
Root-Specific Differential Gene Expression of Selected Highly sDEGs. Example genes identified as highly differentially expressed in shoots and/or roots under hypoxia-salt (**HS**) stress (log2FoldChange). Red line marks the log2FC threshold of log2FC*>*2 and log2FC*<*-2. Gene names along with their corresponding TrinityIDs are provided above each graph. **C**: Control (green); **H**: Hypoxia (yellow); **HS**: Hypoxia-salt (blue); **S**: Salt (red) Abbreviations: SWEET:= Sugar will eventually be exported transporter; SUS:= Sucrose synthase; PCO:= Plant cysteine oxidase; ERFVII:= Ethylene responsive factors group VII; PDC:= Pyruvate decarboxylase; ALDH2:= Aldehyde dehydrogenase; TPP:= Trehalose-6-phosphate phosphatase; VIF2:= Vacuolar/cell wall invertase inhibitor; FBA1:= Fructose-1,6-bisphosphate aldolase 1; G6PDH:= Glucose-6-phosphate dehydrogenase; GAD:= Glutamate decarboxylase; SnRK:= Sucrose non-fermenting 1 related protein kinase

**Figure S11:**
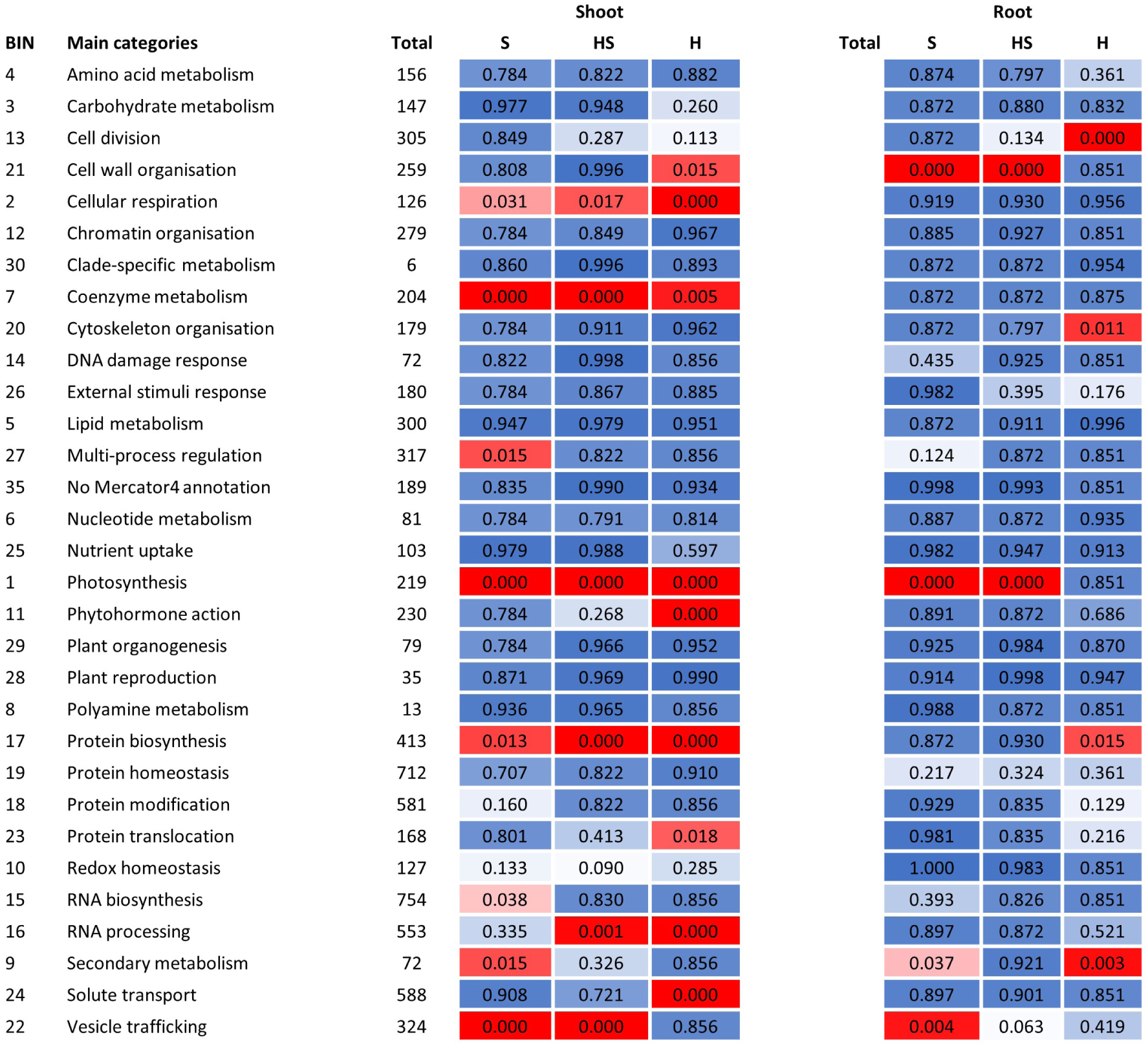
Probability of Enriched Categories in Shoots and Roots. Results of the Wilcoxon-Mann-Whitney test were extracted from MapMan’s statistical tab. Displayed here are BINs, main categories, total element counts within each BIN, and their corresponding probabilities for each category under various conditions in shoot and root data. Enrichment probabilities were calculated according to Wilcoxon-Mann-Whitney; low probabilities are color-coded blue while high probabilities are red. **H**: Hypoxia (yellow); **HS**: Hypoxia-salt (blue); **S**: Salt (red)

**Figure S12:**
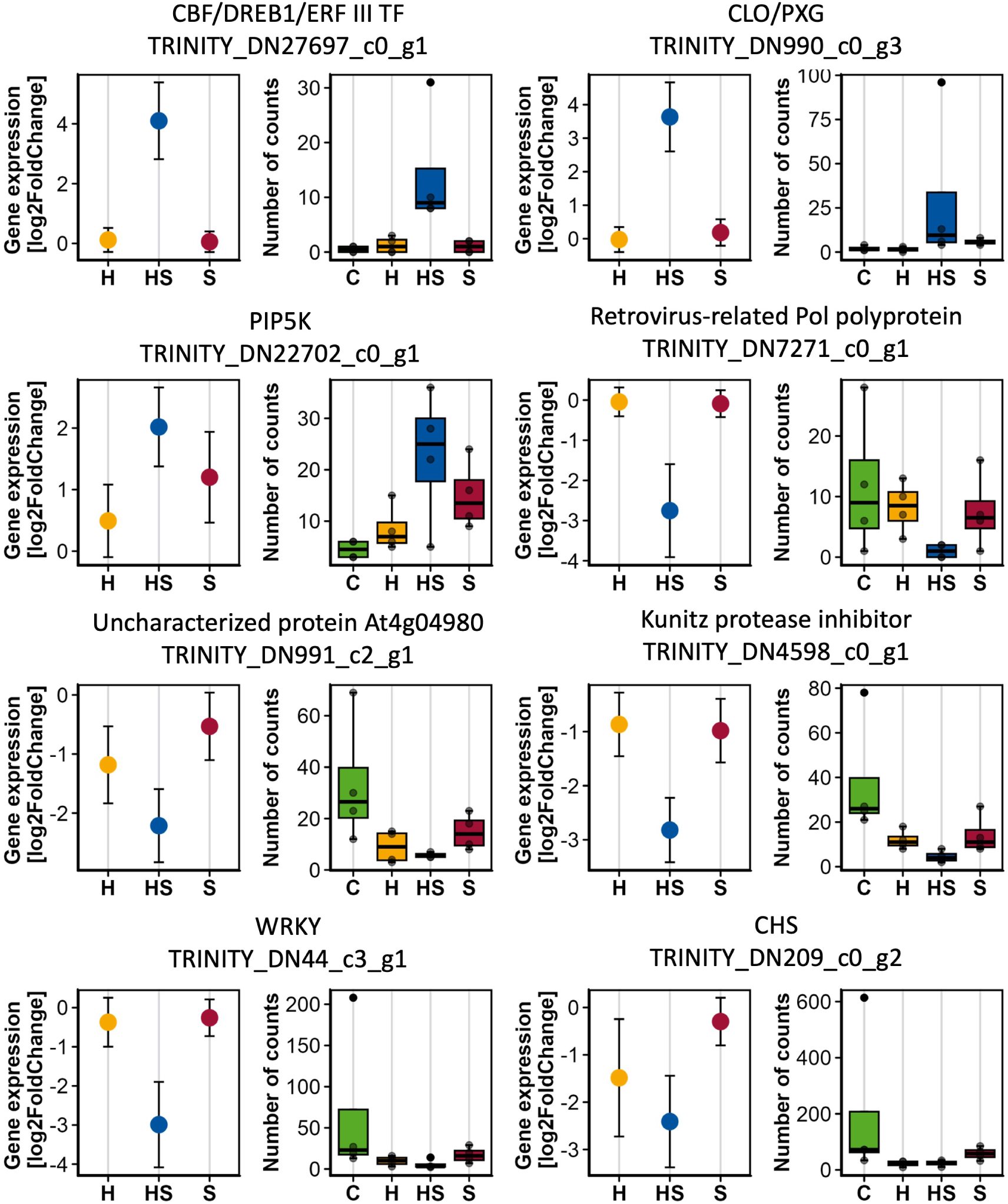
Examples of Unique, Highly Differentially Expressed Synergistic or Antagonistic Hypoxia-Salt Genes of Shoots. Example genes identified as unique, highly differentially expressed, synergistic or antagonistic under hypoxia-salt (**HS**) in shoots. Unique **HS** genes from Venn analysis were filtered for highly gene expression (log2FoldChange*>*2 or log2FoldChange*<*-2) and synergistic and antagonistic characteristics. For each gene, differential gene expression value (log2FoldChange) and standard error were extracted from the DESeq analysis and displayed as dotplot with the respective standard error. The number of counts was extracted from the raw read count table. The four replicate values were displayed for each condition in a boxplot. **C**: Control (green); **H**: Hypoxia (yellow); **HS**: Hypoxia-salt (blue); **S**: Salt (red) Abbreviations: CBF/DREB1/ERFIII TF:= Ethylene responsive factors III transcription factor; CLO/PXG:= Caleosin-type peroxygenase; PIP5K:= Phosphatidylinositol 4-phosphate 5-kinase; WRKY:= WRKY amino acid sequence; CHS:= Chalcone synthase

**Figure S13:**
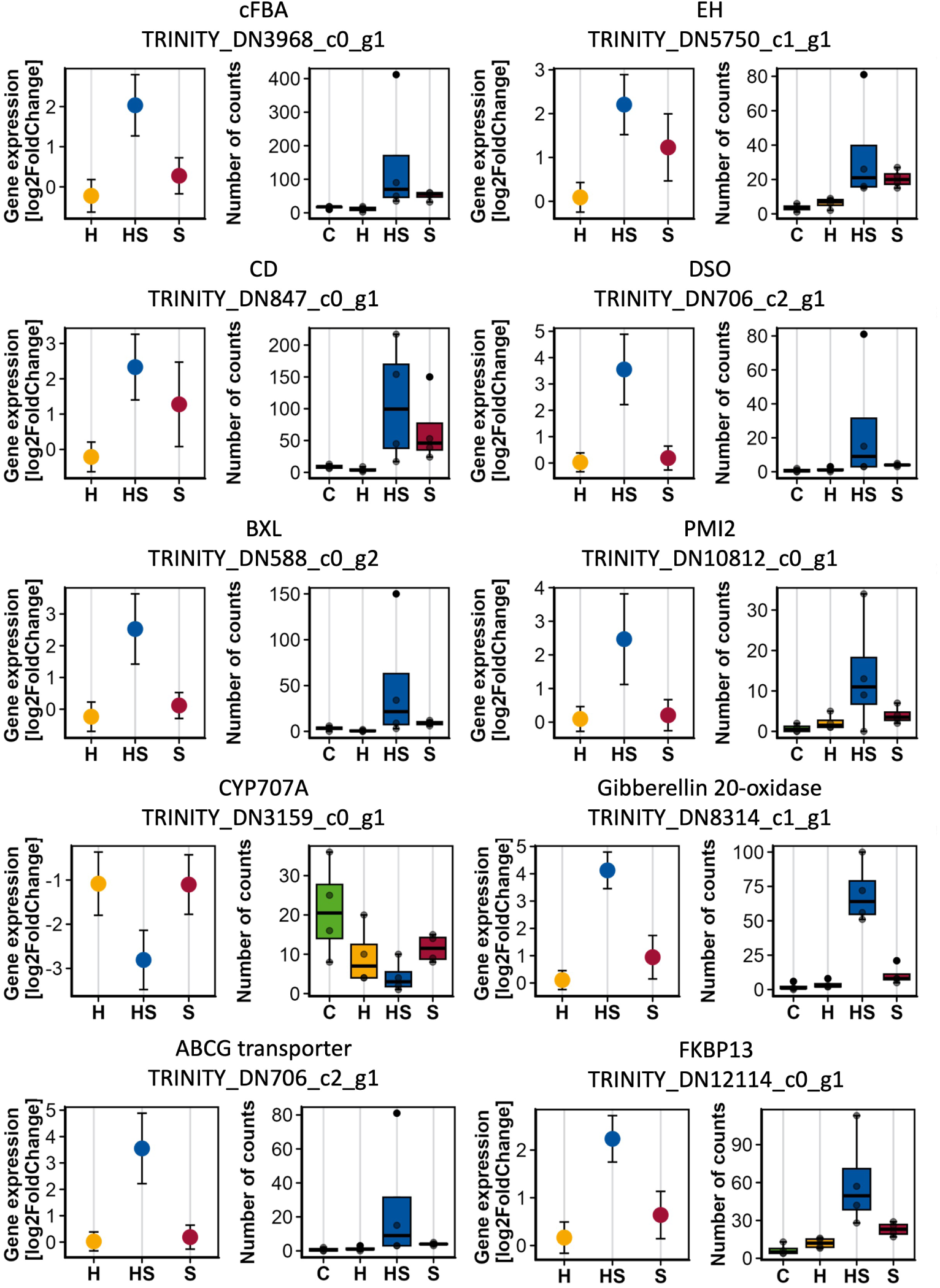
Selection of Unique, Highly Differentially Expressed Synergistic or Antagonistic Hypoxia-Salt Genes of Roots. Example genes identified as unique, highly differentially expressed, synergistic or antagonistic under hypoxia-salt (**HS**) in roots. Unique **HS** genes from Venn analysis were filtered for high gene expression (log2FoldChange*>*2 or log2FoldChange*<*-2) and synergistic and antagonistic characteristics. For each gene, differential gene expression value (log2FoldChange) and standard error were extracted from the DESeq analysis and displayed as dotplot with the respective standard error. The number of counts was extracted from the raw read count table. The four replicate values were displayed for each condition in a boxplot. Differential expression of photosynthesis genes can be explained by either contamination with algae in the hydroponic system, or unusual alteration of gene expression in the roots by light irradiance at the upper sections of the roots due to the cultivation setup. **C**: Control (green); **H**: Hypoxia (yellow); **HS**: Hypoxia-salt (blue); **S**: Salt (red) Abbreviations: FBA:= Fructose-bisphosphate aldolase; EH:= Epoxide hydrolase; CD:= Cutin synthase; DSO:= Suberin/cutin lipid exporter; BXL:= Bifunctional alpha-L-arabinofuranosidase and beta-D-xylosidase; PMI:= WEB1-PMI2 actin filament reorganisation complex; CYP707A:= Abscisic acid 8’-hydroxylase 1; FKBP:= FKBP prolyl isomerase; ABCG:= ATP-binding cassette transporter

**Figure S14:**
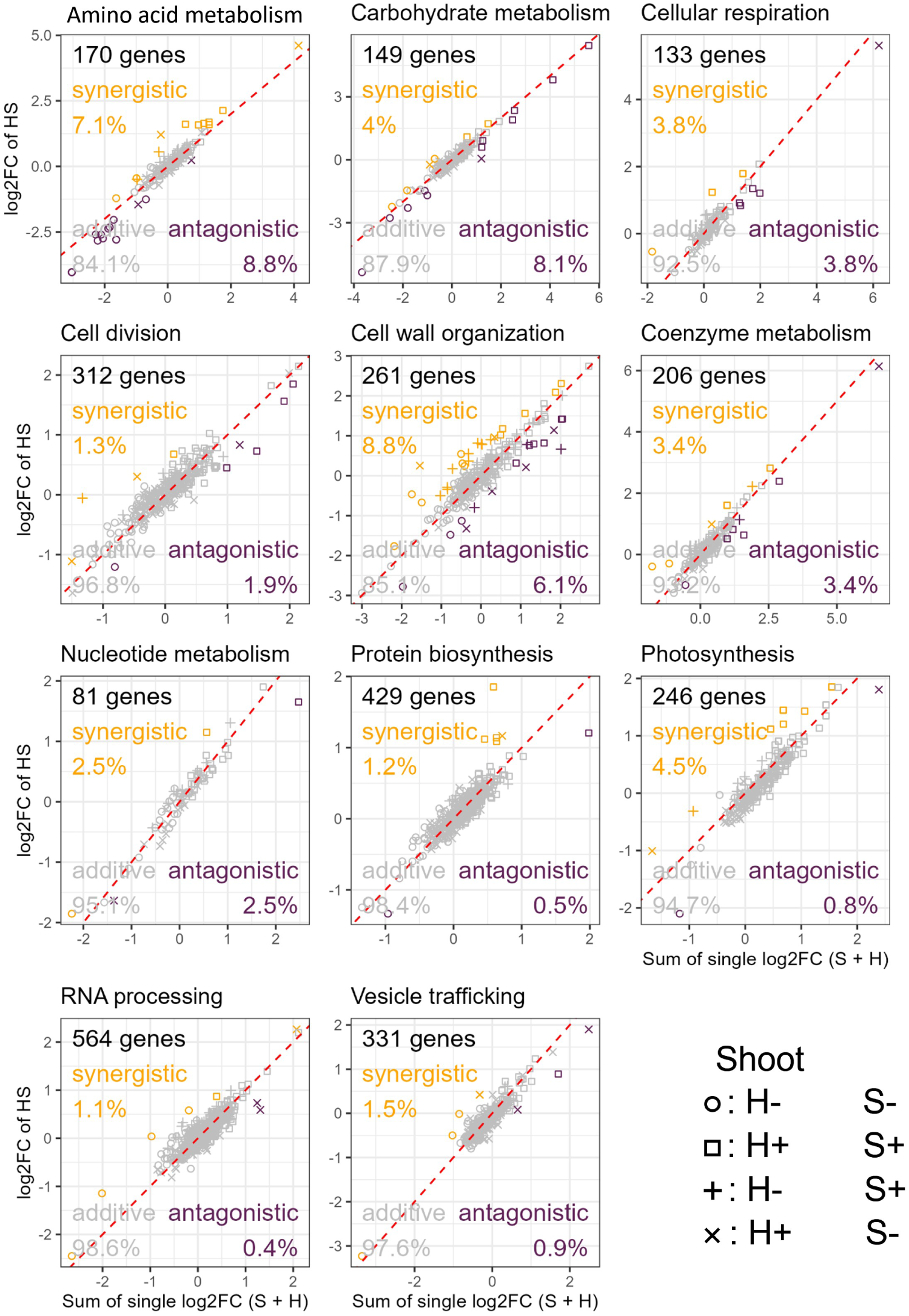
Deviation of Significant Gene Expression Responses from Additive Effects Under Combined Hypoxia-Salt Stress. The relationship between the summed log2FoldChanges (log2FC) of individual stress responses (salt and hypoxia) and the log2FC under simultaneous hypoxia-salt (**HS**) stress in shoots for all enriched functional categories (Suppl. Fig. S11, red). Grey markers denote genes with additive effects, while orange and violet markers indicate genes with synergistic or antagonistic effects, respectively. Additive effects were defined as follows: when the sum of individual stress responses matches the **HS** response (FC(**HS**) = FC(**H**+**S**) ±FC(0.5) confidence interval). Synergistic effects were defined when **HS** expression levels exceed the sum by at least 0.5 FC, and antagonistic effects, when **HS** was below the sum by 0.5 FC threshold. The red diagonal denote the expected trend for additive responses. Symbols represent the sign of log2FC of the individual stress (circle: both (**H** and**S**) negative; square: both (**H** and **S**) positive; +: **H** negative and **S** positive; ×: **H** positive and **S** negative) Abbreviations: FC:= fold change

**Figure S15:**
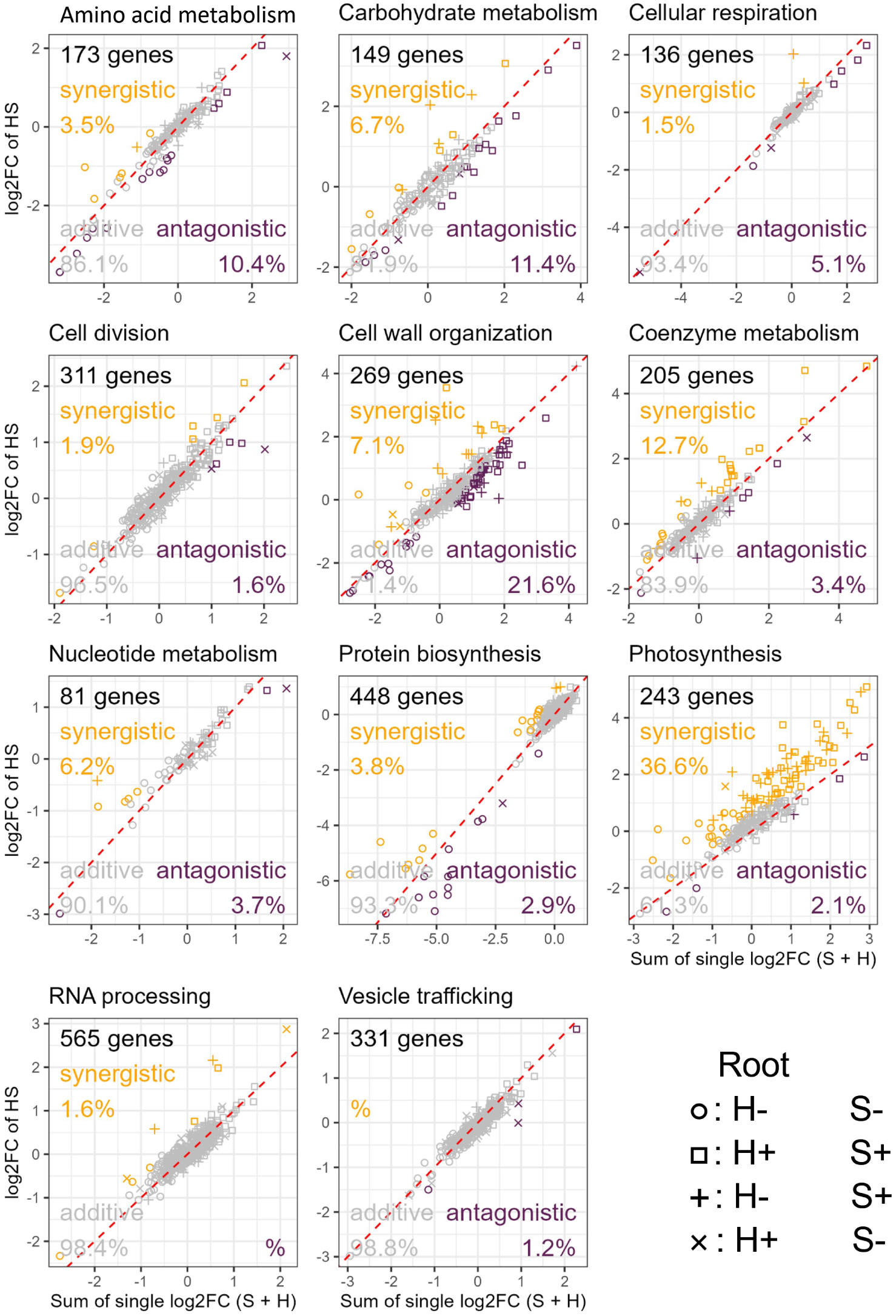
Deviation of Significant Gene Expression Responses from Additive Effects Under Combined Hypoxia-Salt Stress. The relationship between the summed log2FoldChanges (log2FC) of individual stress responses (salt and hypoxia) and the log2FC under simultaneous hypoxia-salt (**HS**) stress in roots for all enriched functional categories (Suppl. Fig. S11, red). Grey markers denote genes with additive effects, while orange and violet markers indicate genes with synergistic or antagonistic effects, respectively. Additive effects were defined as follows: when the sum of individual stress responses matches the **HS** response (FC(**HS**) = FC(**H**+**S**) ±FC(0.5) confidence interval). Synergistic effects were defined when **HS** expression levels exceed the sum by at least 0.5 FC, and antagonistic effects, when **HS** was below the sum by 0.5 FC threshold. The red diagonal denote the expected trend for additive responses. Symbols represent the sign of log2FC of the individual stress (circle: both (**H** and**S**) negative; square: both (**H** and **S**) positive; +: **H** negative and **S** positive; ×: **H** positive and **S** negative). Abbreviations: FC:= fold change

## References

1. Alptekin, B., & Kunkowska, A. B. (2024). How plants adapt to combined and sequential abiotic stresses: A transcriptomics approach. Plant Physiology, 197 (1). 10.1093/plphys/kiaf006

2. Anaconda. (2023). https://anaconda.com

3. Andrews, S. (2010). Fastqc a quality control tool for high throughput sequence data. https://www.bioinformatics.babraham.ac.uk/projects/fastqc/.

4. António, C., Päpke, C., Rocha, M., Diab, H., Limami, A. M., Obata, T., Fernie, A. R., & van Dongen, J. T. (2015). Regulation of primary metabolism in response to low oxygen availability as revealed by carbon and nitrogen isotope redistribution. Plant Physiology, 170(1), 43–56. 10.1104/pp.15.00266

5. Apelt, F., Mavrothalassiti, E., Gupta, S., Machin, F., Olas, J. J., Annunziata, M. G., Schindelasch, D., & Kragler, F. (2021). Shoot and root single cell sequencing reveals tissue- and daytime-specific transcriptome profiles. Plant Physiology, 188(2), 861–878. 10.1093/plphys/kiab537

6. Bailey-Serres, J., Fukao, T., Gibbs, D. J., Holdsworth, M. J., Lee, S. C., Licausi, F., Perata, P., Voesenek, L. A., & van Dongen, J. T. (2012). Making sense of low oxygen sensing. Trends Plant Sci, 17 (3), 129–38. 10.1016/j.tplants.2011.12.004

7. Balfagón, D., Sengupta, S., Gómez-Cadenas, A., Fritschi, F. B., Azad, R. K., Mittler, R., & Zandalinas, S. I. (2019). Jasmonic acid is required for plant acclimation to a combination of high light and heat stress. Plant Physiology, 181(4), 1668–1682. 10.1104/pp.19.00956

8. Barding, J., G. A., Beni, S., Fukao, T., Bailey-Serres, J., & Larive, C. K. (2013). Comparison of gc-ms and nmr for metabolite profiling of rice subjected to submergence stress. J Proteome Res, 12(2), 898–909. 10.1021/pr300953k

9. Baud, S., Vaultier, M.-N., & Rochat, C. (2004). Structure and expression profile of the sucrose synthase multigene family in arabidopsis. Journal of Experimental Botany, 55(396), 397–409. 10.1093/jxb/erh047

10. Behr, J. H., Bouchereau, A., Berardocco, S., Seal, C. E., Flowers, T. J., & Zörb, C. (2017). Metabolic and physiological adjustment of *Suaeda maritima* to combined salinity and hypoxia. Annals of Botany, 119(6), 965–976. 10.1093/aob/mcw282

11. Behr, J. H., Bednarz, H., Gödde, V., Niehaus, K., & Zörb, C. (2021). Metabolic responses of sugar beet to the combined effect of root hypoxia and nacl-salinity. Journal of Plant Physiology, 267, 153545. 10.1016/j.jplph.2021.153545

12. Benjamini, Y., & Hochberg, Y. (2000). On the adaptive control of the false discovery rate in multiple testing with independent statistics. Journal of Educational and Behavioral Statistics, 25(1), 60–83. 10.3102/10769986025001060

13. Billings, W. D., & Mooney, H. A. (1968). The ecology of arctic and alpine plants. Biological Reviews, 43(4), 481–529. 10.1111/j.1469-185x.1968.tb00968.x

14. Bliss, L. C. (1971). Arctic and alpine plant life cycles. Annual review of ecology systematics, 2(1), 405–438.

15. Bolger, A. M., Lohse, M., & Usadel, B. (2014). Trimmomatic: A flexible trimmer for illumina sequence data. Bioinformatics, 30(15), 2114–2120. 10.1093/bioinformatics/btu170

16. Bulut, M., Karakas, E., & Fernie, A. R. (2025). Metabolic responses to multi-stress: An update. Plant Stress, 15, 100729. 10.1016/j.stress.2024.100729

17. Cho, H. Y., Loreti, E., Shih, M. C., & Perata, P. (2021). Energy and sugar signaling during hypoxia. New Phytology, 229(1), 57–63. 10.1111/nph.16326

18. Danecek, P., Bonfield, J. K., Liddle, J., Marshall, J., Ohan, V., Pollard, M. O., Whitwham, A., Keane, T., McCarthy, S. A., Davies, R. M., & Li, H. (2021). Twelve years of samtools and bcftools. Gigascience, 10(2). 10.1093/gigascience/giab008

19. de la Torre, F., El-Azaz, J., Avila, C., & Canovas, F. M. (2013). Deciphering the role of aspartate and prephenate aminotransferase activities in plastid nitrogen metabolism. Plant Physiology, 164(1), 92–104. 10.1104/pp.113.232462

20. Di Martino, C., Delfine, S., Pizzuto, R., Loreto, F., & Fuggi, A. (2003). Free amino acids and glycine betaine in leaf osmoregulation of spinach responding to increasing salt stress. New Phytologist, 158(3), 455–463. 10.1046/j.1469-8137.2003.00770.x

21. Diab, H., & Limami, A. (2016). Reconfiguration of n metabolism upon hypoxia stress and recovery: Roles of alanine aminotransferase (alaat) and glutamate dehydrogenase (gdh). Plants, 5(2), 25. 10.3390/plants5020025

22. Du, L., Li, S., Ding, L., Cheng, X., Kang, Z., & Mao, H. (2022). Genome-wide analysis of trehalose-6-phosphate phosphatases (tpp) gene family in wheat indicates their roles in plant development and stress response. BMC Plant Biology, 22(1). 10.1186/s12870-022-03504-0

23. Ejaz, S., Fahad, S., Anjum, M. A., Nawaz, A., Naz, S., Hussain, S., & Ahmad, S. (2020). Role of osmolytes in the mechanisms of antioxidant defense of plants. In Sustainable agriculture reviews 39 (pp. 95–117). Springer International Publishing. 10.1007/978-3-030-38881-2_4

24. Fürtauer, L., Pschenitschnigg, A., Scharkosi, H., Weckwerth, W., & Nägele, T. (2018). Combined multivariate analysis and machine learning reveals a predictive module of metabolic stress response in *Arabidopsis thaliana*. Molecular Omics, 14(6), 437–449. 10.1039/c8mo00095f

25. Garg, R., Verma, M., Agrawal, S., Shankar, R., Majee, M., & Jain, M. (2013). Deep transcriptome sequencing of wild halophyte rice, *Porteresia coarctata*, provides novel insights into the salinity and submergence tolerance factors. DNA Research, 21(1), 69–84. 10.1093/dnares/dst042

26. Gautam, T., Dutta, M., Jaiswal, V., Zinta, G., Gahlaut, V., & Kumar, S. (2022). Emerging roles of sweet sugar transporters in plant development and abiotic stress responses. Cells, 11(8), 1303. 10.3390/cells11081303

27. Giesbrecht, O., Bonn, C., & Fürtauer, L. (2025). Cytosolic fructose - an underestimated player in the regulation of sucrose biosynthesis. BMC Plant Biology, 25(535). 10.1186/s12870-025-06493-y

28. Glup, G. (1985). Dünen, watt und salzwiesen - schutz und erhaltung von küsten und inseln, tier und pflanzenwelt. Niedersächsischer Minister für Ernährung, Landwirtschaft und Forsten.

29. Gorai, M., Ennajeh, M., Khemira, H., & Neffati, M. (2010). Combined effect of nacl-salinity and hypoxia on growth, photosynthesis, water relations and solute accumulation in *Phragmites australis* plants. *Flora - Morphology, Distribution*, Functional Ecology of Plants, 205(7), 462–470. 10.1016/j.flora.2009.12.021

30. Gutterman, Y. (2012). Seed germination in desert plants. Springer Science & Business Media.

31. Heinemann, B., & Hildebrandt, T. M. (2021). The role of amino acid metabolism in signaling and metabolic adaptation to stress-induced energy deficiency in plants (K.-J. Dietz, Ed.). Journal of Experimental Botany, 72(13), 4634–4645. 10.1093/jxb/erab182

32. Hodge, A., Berta, G., Doussan, C., Merchan, F., & Crespi, M. (2009). Plant root growth, architecture and function. Plant and Soil, 321(1–2), 153–187. 10.1007/s11104-009-9929-9

33. Hsu, F.-C., Chou, M.-Y., Peng, H.-P., Chou, S.-J., & Shih, M.-C. (2011). Insights into hypoxic systemic responses based on analyses of transcriptional regulation in arabidopsis (S.-H. Shiu, Ed.). PLoS ONE, 6(12), e28888. 10.1371/journal.pone.0028888

34. Jacoby, R. P., Taylor, N. L., & Millar, A. H. (2011). The role of mitochondrial respiration in salinity tolerance. Trends in plant science, 16(11), 614–23. 10.1016/j.tplants.2011.08.002

35. Jordine, A., Retzlaff, J., Gens, L., Ehrt, B., Furtauer, L., & van Dongen, J. T. (2024). Introducing the halophyte *Salicornia europaea* to investigate combined impact of salt and tidal submergence conditions. Functional Plant Biology, 51, FP23228. 10.1071/FP23228

36. Kerbler, S. M.-L., Armijos-Jaramillo, V., Lunn, J. E., & Vicente, R. (2023). The trehalose 6-phosphate phosphatase family in plants. Physiologia Plantarum, 175(6). 10.1111/ppl.14096

37. Kim, D., Paggi, J. M., Park, C., Bennett, C., & Salzberg, S. L. (2019). Graph-based genome alignment and genotyping with hisat2 and hisat-genotype. Nature Biotechnology, 37 (8), 907–915. 10.1038/s41587-019-0201-4

38. Liao, Y., Smyth, G. K., & Shi, W. (2014). Featurecounts: An efficient general purpose program for assigning sequence reads to genomic features. Bioinformatics, 30(7), 923–30. 10.1093/bioinformatics/btt656

39. Licausi, F. (2012). Molecular elements of low-oxygen signaling in plants. Physiologia Plantarum, 148(1), 1–8. 10.1111/ppl.12011

40. Licausi, F., Weits, D. A., Pant, B. D., Scheible, W. R., Geigenberger, P., & van Dongen, J. T. (2011). Hypoxia responsive gene expression is mediated by various subsets of transcription factors and mirnas that are determined by the actual oxygen availability. New Phytologist, 190(2), 442–56. 10.1111/j.1469-8137.2010.03451.x

41. Licausi, F., Weits, D. A., Pant, B. D., Scheible, W.-R., Geigenberger, P., & van Dongen, J. T. (2010). Hypoxia responsive gene expression is mediated by various subsets of transcription factors and mirnas that are determined by the actual oxygen availability. New Phytologist, 190(2), 442–456. 10.1111/j.1469-8137.2010.03451.x

42. Liu, F., VanToai, T., Moy, L. P., Bock, G., Linford, L. D., & Quackenbush, J. (2005). Global transcription profiling reveals comprehensive insights into hypoxic response in arabidopsis. Plant Physiology, 137 (3), 1115–29. 10.1104/pp.104.055475

43. Lohse, M., Nagel, A., Herter, T., May, P., Schroda, M., Zrenner, R., Tohge, T., Fernie, A. R., Stitt, M., & Usadel, B. (2013). Mercator: A fast and simple web server for genome scale functional annotation of plant sequence data. Plant, Cell & Environment, 37 (5), 1250–1258. 10.1111/pce.12231

44. Love, M. I., Huber, W., & Anders, S. (2014). Moderated estimation of fold change and dispersion for rna-seq data with deseq2. Genome Biology, 15(12), 550. 10.1186/s13059-014-0550-8

45. Lu, H., Hu, Y., Wang, C., Liu, W., Ma, G., Han, Q., & Ma, D. (2019). Effects of high temperature and drought stress on the expression of gene encoding enzymes and the activity of key enzymes involved in starch biosynthesis in wheat grains. Frontiers in Plant Science, 10. 10.3389/fpls.2019.01414

46. Ma, J., Xiao, X., Li, L., Maggio, A., Zhang, D., Abdelshafy Mohamad, O. A., Van Oosten, M., Huang, G., Sun, Y., Tian, C., & Yao, Y. (2018). Large-scale de novo transcriptome analysis reveals specific gene expression and novel simple sequence repeats markers in salinized roots of the euhalophyte *Salicornia europaea*. Acta Physiologiae Plantarum, 40(8). 10.1007/s11738-018-2702-z

47. Mahalingam, R. (2015). Consideration of combined stress: A crucial paradigm for improving multiple stress tolerance in plants. in ‘combined stress in plants: Physiological, molecular and biochemical aspects. In R. Mahalingam (Ed.), Combined stress in plants: Physiological, molecular and biochemical aspects (pp. 1–26). Springer. https://books.google.de/books?id=Rn65BQAAQBAJ%5C&printsec=frontcover%5C&hl=de%5C&source=gbs_ge_summary_r%5C&cad=0#v=onepage%5C&q%5C&f=true

48. Mahalingam, R., Pandey, P., & Senthil-Kumar, M. (2021). Progress and prospects of concurrent or combined stress studies in plants. Annual Plant Reviews Online, 4(4), 813–868.

49. McCormick, A. J., & Kruger, N. J. (2015). Lack of fructose 2, 6-bisphosphate compromises photosynthesis and growth in arabidopsis in fluctuating environments. The Plant Journal, 81(5), 670–683. 10.1111/tpj.12765

50. Miricescu, A., Brazel, A. J., Beegan, J., Wellmer, F., & Graciet, E. (2023). Transcriptional analysis in multiple barley varieties identifies signatures of waterlogging response. Plant Direct, 7 (8). 10.1002/pld3.518

51. Misra, N., & Gupta, A. K. (2005). Effect of salt stress on proline metabolism in two high yielding genotypes of green gram. Plant Science, 169(2), 331–339. 10.1016/j.plantsci.2005.02.013

52. Mittler, R. (2006). Abiotic stress, the field environment and stress combination. Trends in Plant Science, 11(1), 15–9. 10.1016/j.tplants.2005.11.002

53. Moore, B., Zhou, L., Rolland, F., Hall, Q., Cheng, W.-H., Liu, Y.-X., Hwang, I., Jones, T., & Sheen, J. (2003). Role of the arabidopsis glucose sensor hxk1 in nutrient, light, and hormonal signaling. Science, 300(5617), 332–336. 10.1126/science.1080585

54. Narsai, R., Rocha, M., Geigenberger, P., Whelan, J., & van Dongen, J. T. (2011). Comparative analysis between plant species of transcriptional and metabolic responses to hypoxia. New Phytologist, 190(2), 472–487. 10.1111/j.1469-8137.2010.03589.x

55. Nefissi Ouertani, R., Arasappan, D., Abid, G., Ben Chikha, M., Jardak, R., Mahmoudi, H., Mejri, S., Ghorbel, A., Ruhlman, T. A., & Jansen, R. K. (2021). Transcriptomic analysis of salt-stress-responsive genes in barley roots and leaves. International Journal of Molecular Sciences, 22(15), 8155. 10.3390/ijms22158155

56. O’Leary, B. M., Oh, G. G. K., Lee, C. P., & Millar, A. H. (2019). Metabolite regulatory interactions control plant respiratory metabolism via target of rapamycin (tor) kinase activation. The Plant Cell, 32(3), 666–682. 10.1105/tpc.19.00157

57. Pandey, P., Ramegowda, V., & Senthil-Kumar, M. (2015). Shared and unique responses of plants to multiple individual stresses and stress combinations: Physiological and molecular mechanisms. Frontiers in Plant Science, 6. 10.3389/fpls.2015.00723

58. Patel, M. K., Kumar, M., Li, W., Luo, Y., Burritt, D. J., Alkan, N., & Tran, L.-S. P. (2020). Enhancing salt tolerance of plants: From metabolic reprogramming to exogenous chemical treatments and molecular approaches. Cells, 9(11), 2492. 10.3390/cells9112492

59. Podestá, F. E., & Plaxton, W. C. (1991). Kinetic and regulatory properties of cytosolic pyruvate kinase from germinating castor oil seeds. Biochemical Journal, 279(2), 495–501. 10.1042/bj2790495

60. Posso, D. A., Shimoia, E. P., da-Silva, C. J., Thuy Phan, A. N., Reissig, G. N., da Silva Martins, T., Ehrt, B., Martins, P. D., de Oliveira, A. C. B., Blank, L. M., Borella, J., van Dongen, J. T., & Amarante, L. d. (2025). Soybean tolerance to waterlogging is achieved by detoxifying root lactate via lactate dehydrogenase in leaves and metabolizing malate and succinate. Plant Physiology and Biochemistry, 220, 109520. 10.1016/j.plaphy.2025.109520

61. R Core Team. (2017). R: A language and environment for statistical computing. R Foundation for Statistical Computing. Vienna, Austria. https://www.R-project.org/

62. Ribeiro, C., Stitt, M., & Hotta, C. T. (2022). How stress affects your budget—stress impacts on starch metabolism. Frontiers in Plant Science, 13. 10.3389/fpls.2022.774060

63. Rizhsky, L., Liang, H. J., Shuman, J., Shulaev, V., Davletova, S., & Mittler, R. (2004). When defense pathways collide. the response of arabidopsis to a combination of drought and heat stress. Plant Physiology, 134(4), 1683–1696. 10.1104/pp.103.033431

64. Rosca, M., Mihalache, G., & Stoleru, V. (2023). Tomato responses to salinity stress: From morphological traits to genetic changes. Frontiers in Plant Science, 14. 10.3389/fpls.2023.1118383

65. Safavi-Rizi, V., Herde, M., & Stöhr, C. (2020). Rna-seq reveals novel genes and pathways associated with hypoxia duration and tolerance in tomato root. Scientific Reports, 10(1). 10.1038/s41598-020-57884-0

66. Sarkar, A. K., & Sadhukhan, S. (2022). Imperative role of trehalose metabolism and trehalose-6-phosphate signaling on salt stress responses in plants. Physiologia Plantarum, 174(1). 10.1111/ppl.13647

67. Sasidharan, R., & Mustroph, A. (2011). Plant oxygen sensing is mediated by the n-end rule pathway: A milestone in plant anaerobiosis. The Plant Cell, 23(12), 4173–4183. 10.1105/tpc.111.093880

68. Schwacke, R., Ponce-Soto, G. Y., Krause, K., Bolger, A. M., Arsova, B., Hallab, A., Gruden, K., Stitt, M., Bolger, M. E., & Usadel, B. (2019). Mapman4: A refined protein classification and annotation framework applicable to multi-omics data analysis. Molecular Plant, 12(6), 879–892. 10.1016/j.molp.2019.01.003

69. Sellami, S., Le Hir, R., Thorpe, M. R., Vilaine, F., Wolff, N., Brini, F., & Dinant, S. (2019). Salinity effects on sugar homeostasis and vascular anatomy in the stem of the *Arabidopsis thaliana* inflorescence. International Journal of Molecular Sciences, 20(13), 3167. 10.3390/ijms20133167

70. Sewelam, N., Oshima, Y., Mitsuda, N., & Ohme-Takagi, M. (2014). A step towards understanding plant responses to multiple environmental stresses: A genome-wide study. Plant, cell and environment, 37 (9), 2024–2035. 10.1111/pce.12274

71. Shaar-Moshe, L., Blumwald, E., & Peleg, Z. (2017). Unique physiological and transcriptional shifts under combinations of salinity, drought, and heat. Plant Physiology, 174(1), 421–434. 10.1104/pp.17.00030

72. Shahid, M. A., Sarkhosh, A., Khan, N., Balal, R. M., Ali, S., Rossi, L., Gómez, C., Mattson, N., Nasim, W., & Garcia-Sanchez, F. (2020). Insights into the physiological and biochemical impacts of salt stress on plant growth and development. Agronomy, 10(7), 938. 10.3390/agronomy10070938

73. Shavrukov, Y. (2012). Salt stress or salt shock: Which genes are we studying? Journal of Experimental Botany, 64(1), 119–127. 10.1093/jxb/ers316

74. Sinha, R., Peláez-Vico, M. Á., Shostak, B., Nguyen, T. T., Pascual, L. S., Ogden, A. M., Lyu, Z., Zandalinas, S. I., Joshi, T., Fritschi, F. B., & Mittler, R. (2024). The effects of multifactorial stress combination on rice and maize. Plant Physiology, 194(3), 1358–1369. 10.1093/plphys/kiad557

75. Skorupa, M., Golebiewski, M., Kurnik, K., Niedojadlo, J., Kesy, J., Klamkowski, K., Wojcik, K., Treder, W., Tretyn, A., & Tyburski, J. (2019). Salt stress vs. salt shock - the case of sugar beet and its halophytic ancestor. BMC Plant Biology, 19(1), 57. 10.1186/s12870-019-1661-x

76. Sun, P., Jia, H., Zhang, Y., Li, J., Lu, M., & Hu, J. (2019). Deciphering genetic architecture of adventitious root and related shoot traits in *Populus* using qtl mapping and rna-seq data. International Journal of Molecular Sciences, 20(24), 6114. 10.3390/ijms20246114

77. Sweetlove, L. J., Dunford, R., Ratcliffe, R. G., & Kruger, N. J. (2000). Lactate metabolism in potato tubers deficient in lactate dehydrogenase activity. Plant, Cell & Environment, 23(8), 873–881. 10.1046/j.1365-3040.2000.00605.x

78. Teixeira, J., & Fidalgo, F. (2009). Salt stress affects glutamine synthetase activity and mrna accumulation on potato plants in an organ-dependent manner. Plant Physiology and Biochemistry, 47 (9), 807–813. 10.1016/j.plaphy.2009.05.002

79. van Dongen, J. T., Gupta, K. J., Ramírez-Aguilar, S. J., Araújo, W. L., Nunes-Nesi, A., & Fernie, A. R. (2011). Regulation of respiration in plants: A role for alternative metabolic pathways. Journal of Plant Physiology, 168(12), 1434–43. 10.1016/j.jplph.2010.11.004

80. van Dongen, J. T., Schurr, U., Pfister, M., & Geigenberger, P. (2003). Phloem metabolism and function have to cope with low internal oxygen. Plant Physiology, 131(4), 1529–1543. 10.1104/pp.102.017202

81. van Veen, H., Triozzi, P. M., & Loreti, E. (2024). Metabolic strategies in hypoxic plants. Plant Physiology, 197 (1). 10.1093/plphys/kiae564

82. Vives-Peris, V., López-Climent, M. F., Pérez-Clemente, R. M., & Gómez-Cadenas, A. (2020). Root involvement in plant responses to adverse environmental conditions. Agronomy, 10(7), 942. 10.3390/agronomy10070942

83. Wang, C.-F., Han, G.-L., Yang, Z.-R., Li, Y.-X., & Wang, B.-S. (2022). Plant salinity sensors: Current understanding and future directions. Frontiers in Plant Science, 13. 10.3389/fpls.2022.859224

84. Wang, T., Chen, Y., Zhang, M., Chen, J., Liu, J., Han, H., & Hua, X. (2017). Arabidopsis amino acid permease1 contributes to salt stress-induced proline uptake from exogenous sources. Frontiers in Plant Science, 8. 10.3389/fpls.2017.02182

85. Wang, W., Chen, Q., Xu, S., Liu, W.-C., Zhu, X., & Song, C.-P. (2020). Trehalose-6-phosphate phosphatase e modulates aba-controlled root growth and stomatal movement in arabidopsis. Journal of Integrative Plant Biology, 62(10), 1518–1534. 10.1111/jipb.12925

86. Waters, E. R. (2003). Molecular adaptation and the origin of land plants. Molecular Phylogenetics and Evolution, 29(3), 456–463. 10.1016/j.ympev.2003.07.018

87. Weits, L., Daan A. and Zhou, Giuntoli, B., Carbonare, L. D., Iacopino, S., Piccinini, L., Lombardi, L., Shukla, V., Bui, L. T., Novi, G., van Dongen, J. T., & Licausi, F. (2022). Acquisition of hypoxia inducibility by oxygen sensing n-terminal cysteine oxidase in spermatophytes. Plant Cell Environment. 10.1111/pce.14440

88. Yang, Y., & Guo, Y. (2017). Elucidating the molecular mechanisms mediating plant salt-stress responses. New Phytologist, 217 (2), 523–539. 10.1111/nph.14920

89. Yang, Y., & Guo, Y. (2018). Unraveling salt stress signaling in plants. Journal of Integrative Plant Biology, 60(9), 796–804. 10.1111/jipb.12689

90. Zabalza, A., Van Dongen, J. T., Froehlich, A., Oliver, S. N., Faix, B., Gupta, K. J., Schmälzlin, E., Igal, M., Orcaray, L., Royuela, M., & Geigenberger, P. (2009). Regulation of respiration and fermentation to control the plant internal oxygen concentration. Plant Physiology, 149(2), 1087–1098. 10.1104/pp.108.129288

91. Zandalinas, S. I., & Mittler, R. (2022). Plant responses to multifactorial stress combination. 234(4), 1161–1167. 10.1111/nph.18087

92. Zandalinas, S. I., Sengupta, S., Fritschi, F. B., Azad, R. K., Nechushtai, R., & Mittler, R. (2021). The impact of multifactorial stress combination on plant growth and survival. New Phytologist, 230(3), 1034–1048. 10.1111/nph.17232

93. Zhang, Z., Mao, C., Shi, Z., & Kou, X. (2017). The amino acid metabolic and carbohydrate metabolic pathway play important roles during salt-stress response in tomato. Frontiers in Plant Science, 8. 10.3389/fpls.2017.01231

94. Zheng, C., Wang, Y., Ding, Z., & Zhao, L. (2016). Global transcriptional analysis reveals the complex relationship between tea quality, leaf senescence and the responses to cold-drought combined stress in *Camellia sinensis*. Frontiers in Plant Science, 7. 10.3389/fpls.2016.01858

95. Zhu, A., Ibrahim, J. G., & Love, M. I. (2018). Heavy-tailed prior distributions for sequence count data: Removing the noise and preserving large differences. Bioinformatics, 35(12), 2084–2092. 10.1093/bioinformatics/bty895

